# Mathematical Analysis of Left Ventricular Elastance with respect to Afterload Change During Ejection Phase

**DOI:** 10.1101/2023.09.15.558036

**Authors:** Shiro Kato, Yukiko Himeno, Akira Amano

## Abstract

Since the left ventricle (LV) has pressure (*P*_ιυ_) and volume (*V*_ιυ_), we can define LV elastance from the ratio between *P*_ιυ_ and *V*_ιυ_, termed as “instantaneous elastance.” On the other hand, end-systolic elastance (*E*_*max*_) is known to be a good index of LV contractility, which is measured by the slope of several end-systolic *P*_ιυ_ -*V*_ιυ_ points obtained by using different loads. The word *E*_*max*_ originates from the assumption that LV elastance increases during the ejection phase and attains its maximum at the end-systole. From this concept, we can define another elastance determined by the slopes of isochronous *P*_ιυ_ -*V*_ιυ_ points, that is *P*_ιυ_ -*V*_ιυ_ points at a certain time after the ejection onset time by using different loads. We refer to this elastance as “load-dependent elastance.”

To reveal the relation between these two elastances, we used a hemodynamic model that included a detailed ventricular myocyte contraction model. From the simulation results, we found that the isochronous *P*_ιυ_ -*V*_ιυ_ points lay in one line and that the line slope corresponding to the load-dependent elastance slightly decreased during the ejection phase, which is quite different from the instantaneous elastance.

Subsequently, we analyzed the mechanism determining these elastances from the model equations. We found that instantaneous elastance is directly related to contraction force generated by the ventricular myocyte, but the load-dependent elastance is determined by two factors: one is the transient characteristics of the cardiac cell, i.e., the velocity–dependent force drops characteristics in instantaneous shortening. The other is the force–velocity relationship of the cardiac cell. We also found that the linear isochronous pressure–volume relation is based on the approximately linear relation between the temporal differential of the cellular contraction force and the cellular shortening velocity that results from the combined characteristics of LV and aortic compliances.

## 1. Introduction

*E*_*max*_ is a well-known index of cardiac contractility, which is the slope of the end-systolic pressure-volume relation (ESPVR), where ESPVR is obtained from the end-systolic points of several pressure-volume loops (PV loops) by using different preloads or afterloads on the left ventricle (LV) [43, 42]. The word *E*_*max*_ represents the maximum elastance of LV, and the concept that it represents the contractility of LV is based on the assumption that the LV elastance increases during the ejection phase and reaches its maximum value at the end-systole. *E*_*max*_ is considered a good index of LV contractility as it was considered to be load independent.

There are many researches on *E*_*max*_. For example, several studies have found that *E*_*max*_ increases with adrenergic stimulation [42, 39, 17, 26, 5], and decreases in ailing conditions such as heart failure [1, 3, 6].

However, in recent studies, *E*_*max*_ was found to be afterload-dependent [9] and ESPVR to not be linear but convex [8, 36]. Moreover, it was found that ESPVR’s volume intercept will move with load [11, 7]. These points are briefly summarized in [4].

In these reports, *E*_*max*_ are measured by the slope of pressure and volume relation at end-systole. By generalizing this elastance to the arbitrary time point during ejection phase, we can think of the elastance as the slope of pressure and volume relation at given time, and this elastance is represented as “load-dependent elastance” (*E*_*load*_) in this paper, where several pressure and volume points with different LV loads at a given time are necessary to determine it.

On the other hand, as we can consider LV as a compartment with pressure and volume, from the amount of the pressure change caused by the instantaneous volume change, the physical elastance of LV can be defined by the ratio between the pressure and volume change. This elastance can be recognized as a physical elastance property at a given time. Thus, in this paper this elastance is represented as “instantaneous elastance” (*E*_*inst*_).

Historically, Templeton et al. measured the ratio between the pressure and the volume change by applying the 22 Hz sinusoidal volume change to canine LV [46]. From our definition, we can consider this ratio to be close to *E*_*inst*_. Additionally, they reported that this value was linearly related to pressure throughout the cardiac cycle. However, the temporal change in the ratio was not clearly reported.

Technically, it is very difficult to directly measure LV elastance. There are several reports on the measurement of cardiac tissue elastances [2, 33, 21, 37]. Saeki et al. measured kitten’s papillary muscle stiffness by applying an oscillation of 0.1 to 60 Hz and reported that stiffness changes with the cardiac cycle, but the amplitude of the difference was not large [34].

Historically, instantaneous elastance and load–dependent elastance were conceptually considered to represent similar properties of LV. However, there are very few reports on the measurement of instantaneous elastance; thus, it is difficult to compare these two. We now have cellular contraction models based on molecular level findings capable of reproducing muscle kinetic properties. Therefore, in this paper we try to analyze the characteristics of instantaneous elastance and load–dependent elastance based on the molecular level muscle contraction model by using the hemodynamic model combined with the contraction model.

## 2. Model and Definitions

### 2.1. Model structure

In this research, a simplified hemodynamic model proposed by our group [49, 45] was used for mathematical analysis. The model was constructed from a circulation model, a LV geometry model, and a muscle contraction model.

The variables used in the model equations are summaized in Table 1.

**Table 1.** Variables of the simplified circulation model Variables units meaning.

| Variables | units | meaning |
| --- | --- | --- |
| $t$ | [ms] | simulation time |
| $P_a$ | [mmHg] | aortic pressure |
| $V_a$ | [mL] | aortic volume |
| $P_{lv}$ | [mmHg] | LV pressure |
| $V_{lv}$ | [mL] | LV volume |
| $R_{lv}$ | [cm] | LV internal radius |
| $h_{lv}$ | [cm] | LV wall thickness |
| $L$ | [μm] | half-sarcomere length |
| $F_{ext}$ | [mN/mm<sup>2</sup>] | LV wall tension |
| $F_b$ | [mN/mm<sup>2</sup>] | cellular contraction force |
| $F_p$ | [mN/mm<sup>2</sup>] | passive elastic force |
| $TSCa_3^*$ | [μM] | troponin bound Ca<sup>2+</sup> at power state |
| $TSCa_3^{\sim}$ | [μM] | troponin bound Ca<sup>2+</sup> at weak state |
| $TS^*$ | [μM] | troponin at power state |
| $TS^{\sim}$ | [μM] | troponin at weak state |
| $h_p$ | [μm] | crossbridge length at power state |
| $h_w$ | [μm] | crossbridge length at weak state |
| $X_p$ | [μm] | inextensible portion of L at power state |
| $X_w$ | [μm] | inextensible portion of L at weak state |
| $Q_{rel}$ | [μM/ms] | Ca<sup>2+</sup> release flux |
| $Q_{pump}$ | [μM/ms] | Ca<sup>2+</sup> uptake flux |

### 2.2. Circulation model

Because our aim was to mathematically analyze the pressure and volume relation of the model, we used a simplified circulation model based on the Windkessel model (Figure 1). The parameters in the model were basically imported from the circulation model proposed by Heldt et al. [18] and Liang et al. [25].

**Figure 1:**
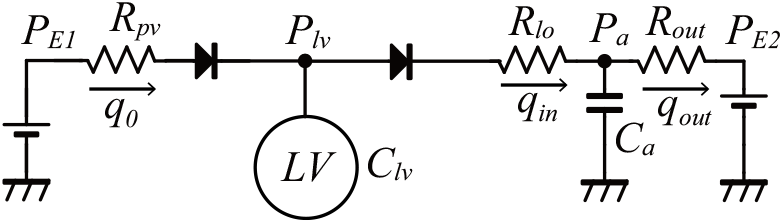
Circulation model

The model included pulmonary venous pressure (*P*_*E*1_), LV pressure (*P*_ιυ_), aortic pressure (*P*_*a*_), and peripheral pressure (*P*_*E*2_) as pressure variables. For simplicity, we used constant values for *P*_*E*1_ and *P*_*E*2_. The model also included pulmonary venous resistance (*R*_*pv*_), aortic resistance (*R*_*lo*_), and peripheral resistance (*R*_*out*_) as vascular resistance parameters. The LV volume was denoted by *V*_ιυ_ and the aortic volume was denoted by *V*_*a*_. The flow between compartments were denoted as *q*_0_ for the LV incoming blood flow, *q*_*in*_ for the aortic blood flow, and *q*_*out*_ for the peripheral blood flow. The aortic compliance was denoted by *C*_*a*_, which has a relation with the aortic pressure and volume as follows.

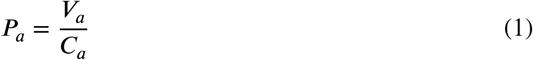

As our aim was to reproduce the baseline hemodynamics, we did not include the baroreflex effect in the model. Moreover, we fixed the cycle length at 1000 [ms]. To evaluate the effect of changes in the afterload, we used several values for *R*_*out*_. The parameters used in the circulation model are shown in Table 2.

**Table 2.** Parameters of the circulation model.

| $P_{E1}$<br>[mmHg] | $P_{E2}$<br>[mmHg] | $R_{pv}$<br>[mmHg · ms/mL] | $R_{lo}$<br>[mmHg · ms/mL] | $R_{out}$<br>[mmHg · ms/mL] | $C_{lv}$<br>[mL/mmHg] | $C_a$<br>[mL/mmHg] |
| --- | --- | --- | --- | --- | --- | --- |
| 10.0 | 15.0 | 27.0 | 2.0 | 1050 - 2050 | variable | 1.5 |

### 2.3. LV geometric model

As our analysis was based on the molecular-level contraction model, a LV geometry model was necessary to relate LV pressure and volume with cellular level contraction force and sarcomere length. We used a measurement-based fitting function that relates 1) the LV internal radius (*R*_ιυ_) to *V*_ιυ_, and 2) *R*_ιυ_ with half-sarcomere length (*L*), respectively. Also, Laplace’s law [27] was used to relate the LV wall tension (*F*_*ext*_) with LV pressure (*P*_ιυ_).

- Relationship between the LV volume and internal radius Utaki et al. [49, 45] defined the equation for the relationship between *R*_ιυ_ and *V*_ιυ_ using the following reported data. Corsi et al. [13] measured the time course of the human LV volume, and Sutton et al. [44] measured the time course of the human LV internal radius. Combining these data, a non-linear relationship between the LV volume and the internal radius was obtained. However, as the resolution of these data was insufficient, they used the time course of the canine LV volume reported by Rodriguez et al. [31] and the time course of the canine LV internal diameter reported by Sabbah et al. [32] to draw a non-linear relationship between the LV volume and internal radius. Given that these are canine data, scaling to human data was performed, and the non-linear equation between the LV volume and internal radius was obtained. In this study, to simplify the mathematical analysis, we approximated the non-linear equation between the LV volume and internal radius into the linear equation as follows.

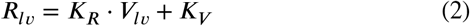

Here, *K*_*R*_ and *K*_*V*_ are constants adjusted to the physiological relation between *R*_ιυ_ and *V*_ιυ_ during end-diastole to end-systole.
- Relationship between the LV myocardial sarcomere length and internal radius Utaki et al. [49, 45] used the following reported data to define the relation between *R*_ιυ_ and *L*. Rodriguez et al. [31] also measured the time course of canine LV sarcomere length. By combining the data with the measured time course of the canine LV internal diameter, as reported by Sabbah et al. [32], a non-linear relationship between the LV volume and sarcomere length was obtained. Also it was assumed that the characteristics were basically similar with canines and humans, thus the scaling factor to the relationship was introduced. In this study, we used the same equation shown in Eq. (3) that Utaki et al. [49] used.

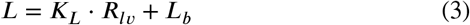

Here, *K*_*L*_ and *L*_*b*_ are constants.
- Simplified LV wall thickness equation LV wall thickness is known to become maximal at the end-systole and minimal at the end-diastole. In a recent report, not only the LV volume but also the LV twist angle was found to be related to the wall thickness [50]. Thus, wall thickness is not always proportional to the LV volume [13, 44], and the quantitative mechanism of wall thickness remains unclear. Utaki et al. [49] assumed that wall thickness is linearly related to the cellular contraction force. However, in this study, to simplify the mathematical analysis, we used constant wall thickness during ejection phase.

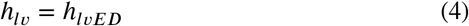

Here, *h*_ιυ_ [cm] denotes the LV wall thickness. Utaki et al. [49] assumed *h*_ιυ_ = 1.00 [cm] at the end-diastole and *h*_ιυ_ = 1.70 [cm] at the end-systole; thus, we used *h*_ιυ*ED*_ = 1.35 [cm] as the constant wall thickness in our analysis.
- Laplace’s law

As Laplace’s law represents well the relation between *P*_ιυ_, *h*_ιυ_, *F*_*ext*_, and *R*_ιυ_ [30], we used this relation in our model.

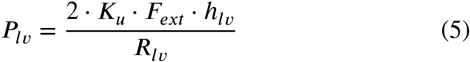

Here, *F*_*ext*_ [mN/mm^2^] denotes the LV wall tension. The calculation method of *F*_*ext*_ is explained in section 2.4. *K*_*u*_ is a parameter that converts units from [mN/mm^2^] to [mmHg].

To reproduce the physiological hemodynamic parameters, we used the model parameter values shown in Table 3.

**Table 3.** Parameters of the LV geometry model.

| $K_R$<br>[cm/mL] | $K_V$<br>[cm] | $K_L$<br>[μm/cm] | $L_b$<br>[μm] | $K_S$ | $K_u$<br>[mmHg/mN/mm<sup>2</sup>] | $h_{lvED}$<br>[cm] |
| --- | --- | --- | --- | --- | --- | --- |
| 0.012 | 1.03 | 0.163 | 0.682 | 7.24 | 7.50064 | 1.35 |

### 2.4. Muscle cell contraction model

In this study, we used a muscle cell contraction model proposed by Negroni and Lascano (NL08 model) [28]. In the model, the cellular contraction force (*F*_*b*_ [mN/mm^2^]) is calculated by multiplying the probability of the power generation states of the troponin system and crossbridge length (*h*_*p*_ [*μ*m], *h*_*w*_ [*μ*m]).

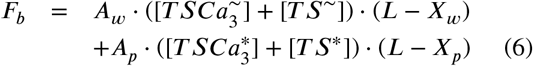

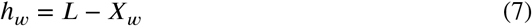

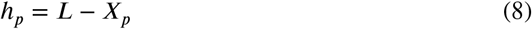

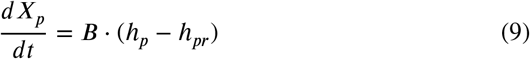

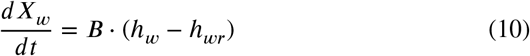

*A*_w_ and *A*_p_ represent the spring rate of crossbridges in the weak (w) and power (p) states that generate *F*_*b*_. 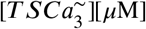, [*TS*∽] [*μ*M], [*TS*Ca^*^] [*μ*M] and [*TS*^*^] [*μ*M] represent the concentrations of th^3^e troponin system forming the crossbridges in the weak (∽) and the power (*) states. *X* [*μ*m] is the non-elastic portion of the contractile element and *L* - *X*_w_ and *L* - *X*_p_ are extensions of the attached crossbridges in the weak and power states, abbreviated as *h*_*w*_ and *h*_*p*_ (Eqs. (7), (8)), respectively. The temporal differential of *X*_p_ and *X*_w_ are given by the products of the crossbridge attaching ratio (*B*) and the crossbridge extension (*h*_*p*_ -h_pr_, *h*_*w*_ -h_wr_), which are different from the initial length(*h*_*p*r_[*μ*m], *h*_*w*r_[*μ*m]) shown in Eqs. (9) and (10). Note that [*TS*Ca^∽^], [*TS*^∽^], [*TS*Ca^*^], [*TS*^*^], *h*_*p*_, and *h*_*w*_ are calculated using the equations in the original paper of the NL08 model [28] with modifications of g and g_*d*_ to be explained later.

As the characteristics of the end-diastolic pressure volume relationship (EDPVR) are similar in rats [22] and humans [12], by linearly scaling the force axis with the identical half-sarcomere length axis, we used the following mammalian exponential function as a human passive elastic force (*F*_*p*_ [mN/mm^2^]) model showing good agreement with the experimental data [48, 16]. The form of this equation was based on the equation used by Shim et al. [38] and Landesberg et al. [23].

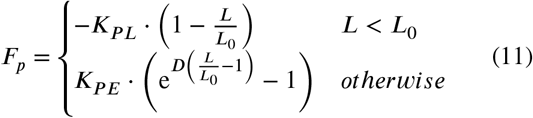

Note that *L*_0_ [*μ*m] is resting half sarcomere length. *D, K*_*PL*_ [mN/mm^2^], and *K*_*PE*_ [mN/mm^2^] are constant parameters that determine the properties of the passive elastic component. The parameter values were manually adjusted to reproduce physiological human hemodynamics (Table 4).

**Table 4.** Parameters in Eq. (11).

| $K_{PE}$ $[\text{mN}/\text{mm}^2]$ | $K_{PL}$ $[\text{mN}/\text{mm}^2]$ | $D$ | $L_0$ $[\mu\text{m}]$ |
| --- | --- | --- | --- |
| 0.2692 | 7.561 | 28.08 | 1.043 |

Since *F*_*p*_ is usually measured using a piece of tissue or an LV cavity we can consider that the characteristics of *F*_*p*_ are compatible with the macroscopic properties. On the other hand, since *F*_*b*_ is usually measured with a single cell or a small piece of ventricular fiber in which the effective cross-sectional area is difficult to measure, the measured force may contain large scale errors. We thus introduced a scale factor, *K*_*s*_, which is multiplied only with *F*_*b*_ to adjust the cellular contraction force. *K*_*s*_ was determined using the method proposed by Utaki et al. [49], which resulted in *K*_*s*_=7.24. Finally, LV wall tension *F*_*ext*_ [mN/mm^2^] in Eq. (5) was cal-culated as follows.

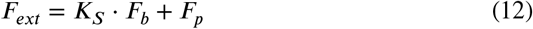

The contraction time course is controlled by *Ca*^2+^, and *Ca*^2+^ release is controlled by the *Ca*^2+^ release equation. The release and absorption of *Ca*^2+^ by the sarcoplasmic reticulum (*Q*_r*el*_ [*μ*M/ms] and *Q*_pu*mp*_ [*μ*M/ms]) in the NL08 model are expressed by the following equations.

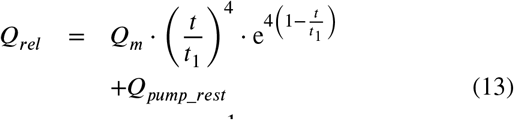

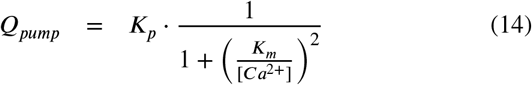

Note that, *t* [ms] is the time parameter, [*Ca*^2+^] [*μ*M] is the concentration of *Ca*^2+^, *Q*_*m*_ [*μ*M/ms] is the maximum level of *Ca*^2+^ release, *t*_1_ [ms] is the interval to maximum *Q*_r*el*_, *Q*_*pum*p_r*est*_ [*μ*M/ms] determines [*Ca*^2+^] at rest, *K*_*p*_ [*μ*M/ms] is the maximum value of *Q*_pu*mp*_, and *K*_*m*_ [*μ*M] the value of [*Ca*^2+^] for *Q*_pu*mp*_ = *K*_*p*_⁄2. Parameter values used in Eq. (13) and (14) are shown in Table 5.

**Table 5.** Parameters used in Eq. (13) and (14).

| $Q_m$<br>[$\mu\text{M}/\text{ms}$] | $t_1$<br>[ms] | $Q_{pump\_rest}$<br>[$\mu\text{M}/\text{ms}$] | $K_p$<br>[$\mu\text{M}/\text{ms}$] | $K_m$<br>[$\mu\text{M}$] |
| --- | --- | --- | --- | --- |
| 3.2 | 8 | 0.03 | 0.15 | 0.2 |

The NL08 model is known to have a problem in the filling phase, where *F*_*b*_ rapidly decreases when *L* extends. To improve the filling phase characteristics, *K*_*γ*_ was introduced in [49], and we used this modification in our model as follows.

**Table 6.** Parameters used to calculate cellular contraction force (*F*_*b*_)

|  |  |  |
| --- | --- | --- |
| $A_p$ | 2700 | [mN mm<sup>-2</sup> μm<sup>-1</sup> μM<sup>-1</sup>] |
| $A_w$ | 540 | [mN mm<sup>-2</sup> μm<sup>-1</sup> μM<sup>-1</sup>] |
| $h_{pr}$ | 0.006 | [μm] |
| $h_{wr}$ | 0.0001 | [μm] |
| $B$ | 0.5 | [ms<sup>-1</sup>] |
| $Y_c$ | 1.0 | [ms<sup>-1</sup> μm<sup>-2</sup>] |
| $Y_d$ | 0.0333 | [ms<sup>-1</sup>] |
| $Y_v$ | 1.5 | [ms<sup>-1</sup>] |
| $Y_{vd}$ | 1.5 | [ms<sup>-1</sup>] |
| $Z_a$ | 0.0023 | [ms<sup>-1</sup>] |
| $L_c$ | 1.2 | [μm] |
| $K_\gamma$ | 85.3 | |

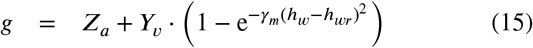

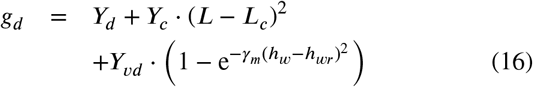

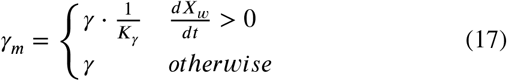

Note that, *h*_*w*r_ [*μ*m] is the steady state extension of the attached crossbridges in the weak state. *Z*_a_ [1/ms] and *Y*_*d*_ [1/ms] are crossbridge dissociation constants, *Y*_*v*_ [1/ms] and *Y*_*vd*_ [1/ms] are model parameters for the weakly-attached crossbridge extension, and *Y*_*c*_ [1/ms/*μ*m^2^], and *L*_*c*_ [*μ*m] are model parameters related to the half-sarcomere length.

### 2.5. Mathematical definitions of elastance

- instantaneous elastance Considering LV as one elastic compartment, the instantaneous elastance *E*_*inst*_ can be defined by the ratio between the instantaneous changes in pressure (*P*_ιυ_) and small changes in volume (*V*_ιυ_), as can be defined as follows.

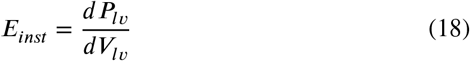
- load-dependent elastance For the heart, pressure and the volume follow a different pressure-volume curve if the afterload is different. Usually, the end-systolic pressure volume curve is measured by measuring several pressure-volume curves with different loads; thus, we can define load-dependent elastance *E*_*load*_ by the ratio between the changes in pressure with changes in afterload 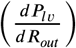 and volume changes with the changes in the afterload 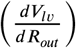 as follows.

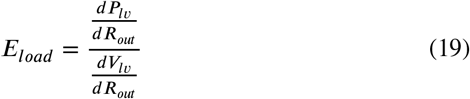

## 3. Simulation

### 3.1 Simulation conditions

Simulations were performed with the *simBio* system [35] in 0.01 [ms] intervals until a steady state was reached, requiring around 100 cardiac cycles. In this study, as a simulation condition, *P*_*a*_ and *V*_*a*_ at the onset time of ejection were fixed to constant values of 70.0 [mmHg] and 105.0 [mL], respectively. In addition, *P*_ιυ_ and *V*_ιυ_ at the onset time of ejection were fixed to 70.0 [mmHg] and 126.419 [mL], respectively. We also measured the *E*_*inst*_ by decreasing the *V*_ιυ_ by 0.25 % (Δ*V*_ιυ_) at a certain time in the ejection phase and by evaluating the pressure drop (Δ*P*_ιυ_) in the next time interval. Thus, *E*_*inst*_ was calculated by Δ*P*_ιυ_⁄Δ*V*_ιυ_.

### 3.2 Simulation results

To evaluate the influence of the afterload on pressure (*P*_ιυ_) and volume (*V*_ιυ_), we performed a simulation with several peripheral resistance values: *R*_*out*_ = 1050, 1200, 1400, 1650, and 2050 [mmHg *⋅* ms/mL]. Under these conditions, the PV loops are shown in Figure 2 and the enlarged ejection phase in Figure 3. In both figures, the isochronous points of *P*_ιυ_ -*V*_ιυ_ are shown at 50, 100, 150, 200, 250, and 280 [ms] after the onset time of the ejection, and the isochronous linear approximation lines of each time point are also shown in these figures. The values of each slope (*E*_*load*_), *V*_ιυ_-axis intercepts (*V*_0_), and *R*^2^ of these lines are shown in Table 7. From the results, the isochronous lines had high *R*^2^ values, thus the isochronous points of *P*_ιυ_ -*V*_ιυ_ had high linearity during the ejection phase. We also noted that *V*_0_ greatly decreased with time, but after the initial drop, *E*_*load*_ slightly decreased over time, although the amount of decrease was quite small as we can see in Figure 3. With the instantaneous volume drop simulation, we also measured *E*_*inst*_ at the same time points shown in Table 7. Notably, *E*_*inst*_ slightly increased and decreased during the ejection phase.

**Table 7.** Changes in *E*_*inst*_, *E*_*load*_, *V*_0_ with time, and its *R*^2^.

| $t$ [ms] | $E_{inst}$<br>($R_{out} = 1050$) | $E_{load}$ | $V_0$ | $R^2$ |
| --- | --- | --- | --- | --- |
| 50 | 49.53 | 10.6280 | 98.2267 | 0.999997 |
| 100 | 52.62 | 7.8969 | 76.7246 | 0.999989 |
| 150 | 54.04 | 6.7593 | 58.9819 | 0.999977 |
| 200 | 53.83 | 6.1503 | 44.9864 | 0.999963 |
| 250 | 51.37 | 5.7014 | 34.8865 | 0.999956 |
| 280 | 47.78 | 5.3194 | 30.8502 | 0.999953 |

**Figure 2:**
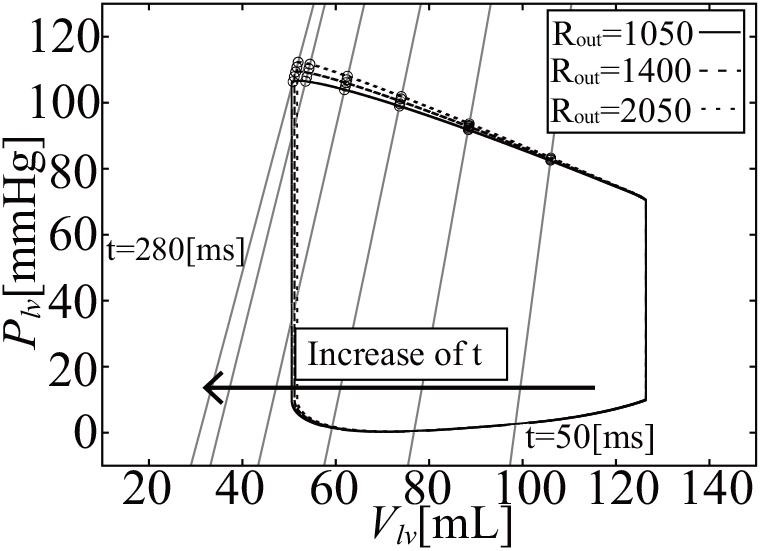
PVloops and isochronous *P*_ιυ_ -*V*_ιυ_ relations

**Figure 3:**
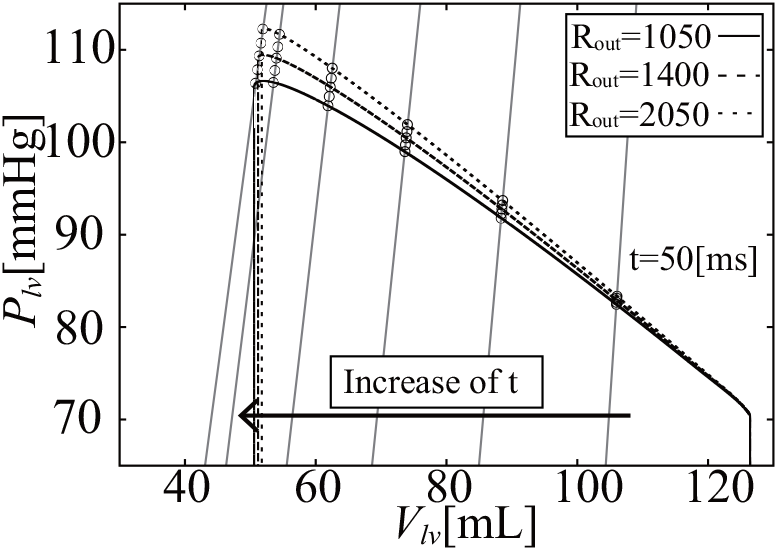
Isochronous *P*_*l,v*_ -*V*_*l,v*_ relations during the ejection phase

In the model, the relation between volume (*V*_*l,v*_) and half-sarcomere length (*L*) is linear, as shown in Eqs. (2), (3). On the other hand, although the relation between LV pressure (*P*_*l,v*_) and wall tension (*F*_*ext*_) is nonlinear, as shown in Eq. (5), it can be approximated as linear for a small change of *P*_*l,v*_. Thus, if we convert the isochronous relation of *P*_*l,v*_ -*V*_*l,v*_ in Figure 3 into the isochronous relation of *F*_*ext*_ -*L* shown in Figure 4, the relation remains linear. The linear approximations for the isochronous points of *F*_*ext*_ -*L* have high-linearity, as evidenced by *R*^2^ in Table 8. The slopes (*k*_a_) of these lines slightly decreased, but they appear to be almost parallel.

**Table 8.** Changes in the slope of the isochronous *F*_*ext*_ -*L* relation (*k*_a_) with time and its *R*^2^.

| $t$ [ms] | $k_a$ | $R^2$ |
| --- | --- | --- |
| 50 | 642.82 | 0.999997 |
| 100 | 445.07 | 0.999988 |
| 150 | 357.38 | 0.999974 |
| 200 | 307.93 | 0.999956 |
| 250 | 274.58 | 0.999939 |
| 280 | 254.25 | 0.999936 |

**Figure 4:**
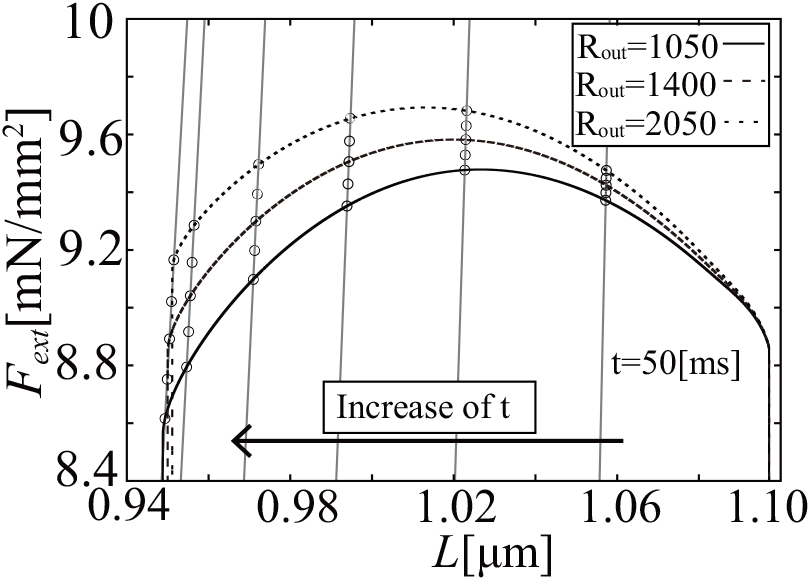
Isochronous *F*_*ext*_ -*L* relation during the ejection phase

## 4. Analysis

### 4.1 Elastance of simplified hemodynamic model

From the simulation results, it can be seen that *E*_*inst*_ slightly increased and then decreased, and its time course was similar to that of [*Ca*^2+^]. On the other hand, *E*_*load*_ was markedly different from *E*_*inst*_ at each time point. *E*_*load*_ initially decreased by around 20 %, and decreased slightly over time. On the other hand, at a certain time point, both elastances were independent of the afterload (*R*_*out*_). The mathematical reasons for these findings are discussed in the following section.

### 4.2 *E*_*inst*_ in the simplified hemodynamic model

*E*_*inst*_ was mathematically defined by Eq. (18), and by using this model, we can represent *E*_*inst*_ as follows.

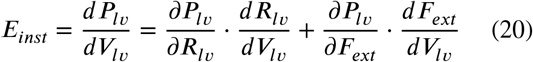

From Eq. (5), the partial differential 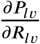 can be derived as follows.

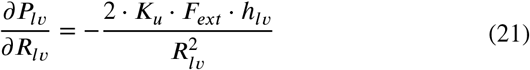

Differential 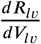 can be derived from Eq. (2) as follows.

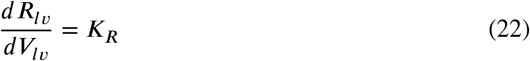

Similarly, from Eq. (5), the partial differential 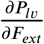 can be derived as follows.

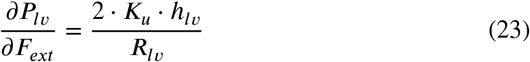

From Eqs. (2), (3), and (12), the differential 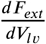 can be derived as follows.

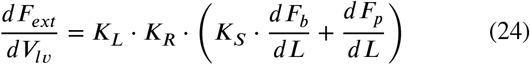

The elastance of the cellular contraction force term 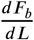 can be derived by differentiating Eq. (6) with *L* as follow

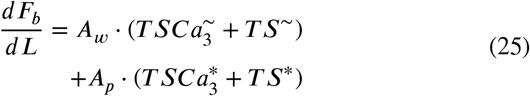

Also, by differentiating Eq. (11) with *L*,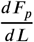 can be derived as follows.

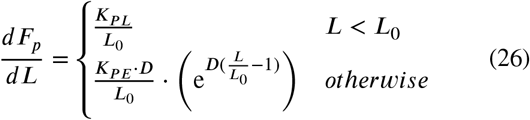

Although Eqs. (25) and (26) appear in Eq. (24), from the simulation results, 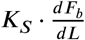 was about 100 times larger than 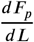 Thus, we can neglect the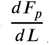 term in Eq. (24).

In this model, *E*_*inst*_ is denoted by Eq. (20). From the simulation results, we found that the second term of the right-hand side of Eq. (20) was dominant. The second term of the right-hand side of Eq. (20) is the product of Eq. (23) and Eq. (24), that represents the changes in the LV pressure by the changes in the LV wall tension, and the changes in the LV wall tension by the changes in the LV volume, respectively. Since 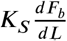 term was dominant in Eq. (24), *E*_*inst*_ can be considered to be proportional to the amount of the attached crossbridges as shown in Eq. (25). And, since the attached crossbridges are formed by the increase in *Ca*^2+^, *E*_*inst*_ tends to follow changes in *Ca*^2+^ concentration. Thus, *E*_*inst*_ can be understood as the LV wall tension elastance mainly determined by the *Ca*^2+^ concentration.

### 4.3 E_loa*d*_ in the simplified hemodynamic model

*E*_*load*_ was mathematically defined as Eq. (19). Although the simulation conditions were not physiological, we fixed *P*_*l,v*_ at the onset time of ejection to simplify the mathematical analysis. The phenomena in the ejection phase can be divided into two steps: 1) initial phase and 2) steady state transition during the ejection phase.

step 1 Immediately after the onset time of ejection, transient variations in crossbridge extension changes occur in short time periods, depending on the afterload.

step 2 The linear isochrones of the *P*_*l,v*_ -*V*_*l,v*_ relations remain linear during the ejection phase with small changes in their slopes.

#### 4.3.1 step 1

It is known that the response to transient force is observed in the initial phase of isovelocity-shortening experiments using skeletal muscles. However, the phenomenon is difficult to measure under accurate isovelocity conditions in real experiments [20, 47, 15, 19]. At this phase, the contraction force changes significantly depending on the velocity. In our analysis, as *P*_*l,v*_ at the onset time of ejection was fixed, the shortening velocity of the half-sarcomere length 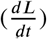 becomes roughly proportional to the inverse of the afterload (*R*_*out*_), corresponding to the quasi isovelocity-shortening condition. Thus, we can predict that the variation of the initial cellular contraction force resulting from the variations of the afterload is caused by the same mechanism.

We performed an isovelocity-shortening simulation by using the NL08 model under the condition of the onset time of ejection. As [*Ca*^2+^] can be considered as constant for this short period, even if the afterload (*R*_*out*_) is different, we fixed [*Ca*^2+^] as 0.3[*μ*M], and performed an isovelocity-shortening simulation of the half-sarcomere length 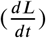 with shortening velocities of 0.1, 0.15, and 0.20 [*μ*m/^*d*^s^*t*^] corresponding to the condition close to the hemodynamics simulation afterloads. The time courses of *L, h*_*p*_, and *F*_*b*_ are shown in Figures 5, 6, and 7, respectively. Note that the isovelocity condition starts at *t* = 100 [ms] in these figures.

**Figure 5:**
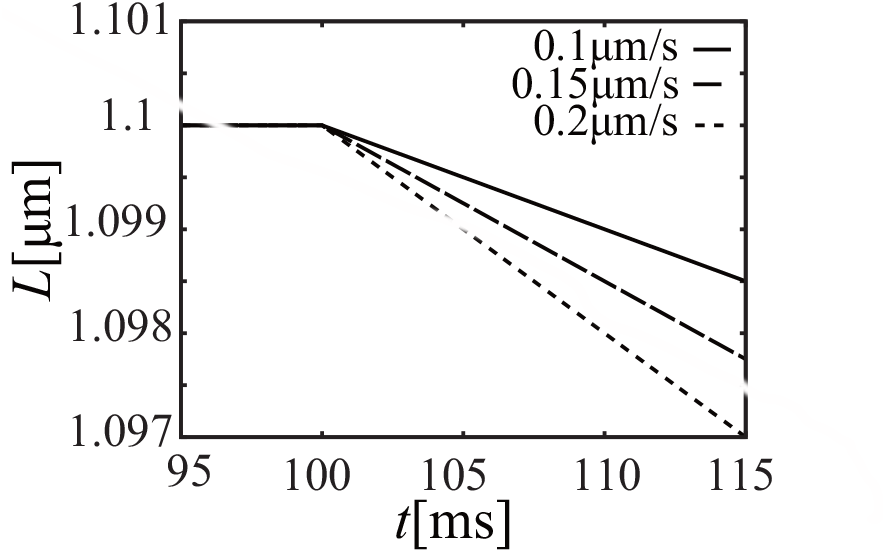
Time course of half-sarcomere length (*L*)

**Figure 6:**
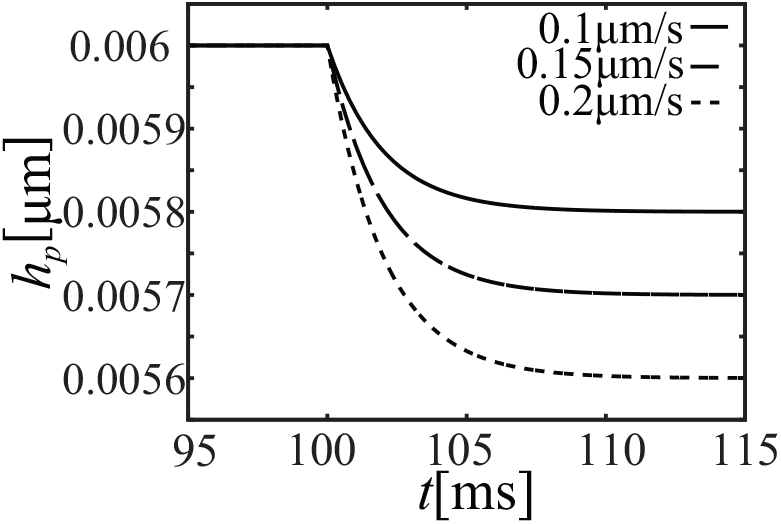
Time course of the crossbridge length (*h*_*p*_)

**Figure 7:**
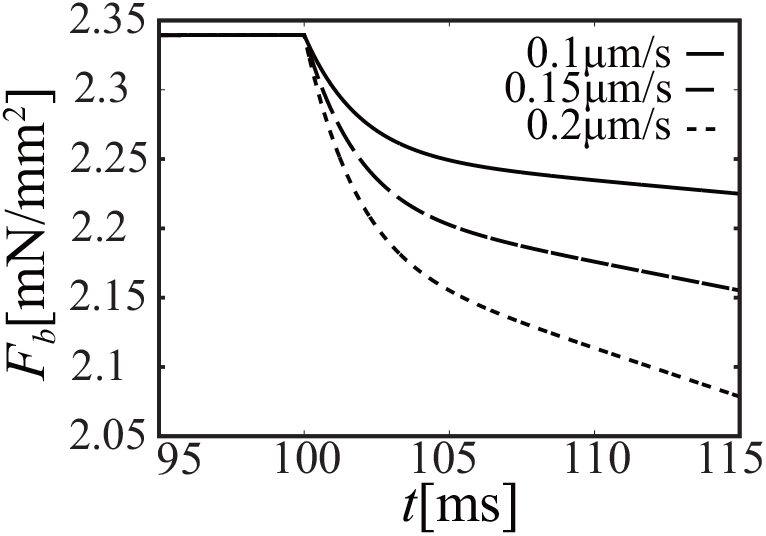
Time course of the cellular contraction force (*F*_*b*_)

As shown in Figures 6 and 7, crossbridge length (*h*_*p*_) of the power state becomes constant after an initial decrease and the time course of *F*_*b*_ is similar to that of *h*_*p*_.

From the model equations, the shortening velocity of the half-sarcomere length 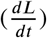 was found to be inversely proportional to the afterload (*R*_*out*_). Thus, *h*_*p*_ changes depending on Eqs. (8), (9) as follows.

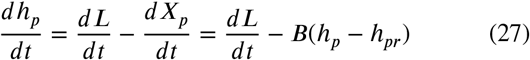

After the initial decrease of *h*_*p*_, it exponentially got close to the steady state length *h*_*p,ss*_, which can be calculated from 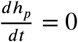 as follows.

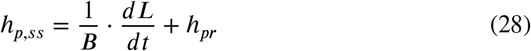

From this equation, *h*_*p*_ decreased from the resting cross-bridge length (*h*_*p*_) to the steady state length that is linearly related with the shortening velocity of the half-sarcomere length 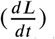 This is what produces the well-known characteristics of the force-velocity relationship. Thus, if the troponin system probabilities are constant, the cellular contraction force (*F*_*b*_) is proportional to *h*_*p*_, which is also proportional to 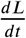 As a result, the isochronous *F*_*ext*_ -*L* (≈ F_b_ -*L*) points located at the same position of the onset time of ejection, move in the *L* direction, depending on the velocity 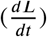 determined by the afterload. However, the movements don e in the *F*_*b*_ direction are maintained by the relation between 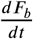 and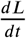, which can be approximated with a linear relation as shown in Eq. (27). Thus, the initial cellular contraction force and length relation becomes almost linear.

In the simulation results in section 3.2, the shortening elocity of the half-sarcomere length 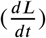 was affected by the cellular contraction force (*F*_*b*_); thus, its ratio was slightly different from the inverse of the afterload (*R*_*out*_). However, the difference between *h*_*p*_ and *h*_*p,ss*_ was around 2.3 % at 15 [ms] after the onset time of ejection.

#### 4.3.2 step 2

As explained in the previous section, the fixed *F*_*ext*_ -*L* points with different afterloads (i.e. fixed *P*_*l,v*_ -*V*_*l,v*_ points) proceeded to isochrones immediately after the onset time of ejection.

Assuming that the relation between *F*_*ext*_ and *L* at a certain time *t*_0_ during the ejection phase became linear, then the following equation holds.

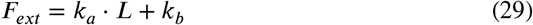

If we assume that linearity is maintained after the small period (Δ*t*), then the following equation holds.

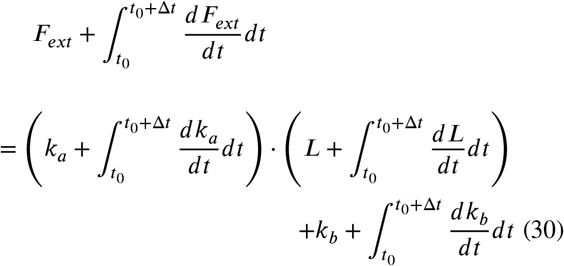

In the differential form, the following equation holds.

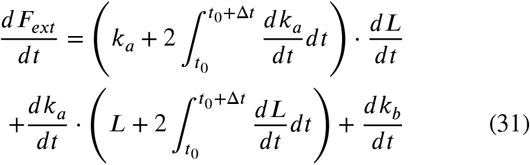

Here, we introduce the assumption that changes in the slope of *k*_a_ from the *F*_*ext*_ -*L* relation is small and can be neglected in a short time period, i.e., 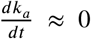 then, the following equation holds.

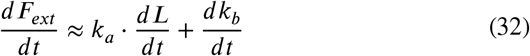

Thus, if the model equations can be decomposed into the above equation form, we can say that the model itself has in its nature the property of a linear *F*_*ext*_ -*L* relation.

First, by temporally differentiating the LV wall tension (Eq. (12)), we get 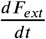 as follows.

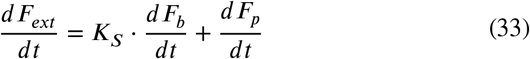

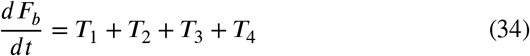

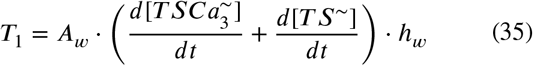

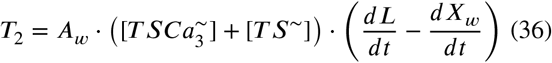

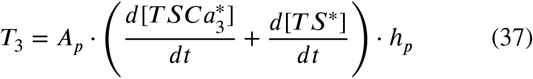

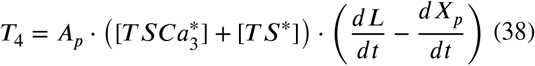

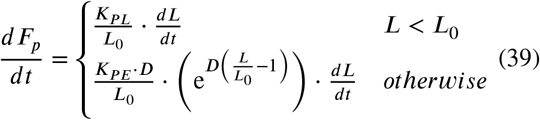

In Eq. (39), if *L L*_0_, then 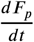 is proportional to 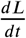.

If L ≥ *L*_0_, then 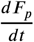 is a function of *L* and 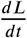 However, from the simulation results, changes in *L* with *R*_*out*_ is 0.63%. Thus, we approximated 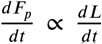, and the passive elastic force component is approximated by using Eq. (32).

On the other hand, in Eq. (34), 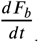 is composed of four terms. If we look at these terms, *T*_1_ and *T*_2_ are similar to *T*_3_ and *T*_4_, respectively; thus, the mathematical analysis of *T*_1_ and *T*_2_ becomes the same for *T*_3_ and *T*_4_, respectively. Moreover, *A*_p_ is five times larger than *A*_w_. Therefore, we can reduce our analysis of Eq. (34) to the analysis of *T*_3_ and *T*_4_.

*T*_3_ is a product of the crossbridge length (*h*_*p*_) and the temporal differential term of 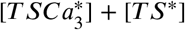. As explained in section 4.3.1, *h* decreases fro^3^m the resting crossbridge length (*h*_pr_), and the amount of decrease is proportional to the shortening velocity of the half-sarcomere length 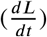. However, as we can see from Figure 6, the decrease of is s small that it can be approximated by *h*_pr_ In addition, the temporal differential of 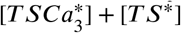 is affected by *L* and 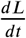 however, the effect is not significant. In general, *h*_*p*_ can be approximated to *h*_*p*r_ and then *F*_*b*_ becomes proportional to 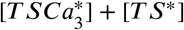 As the temporal differential of 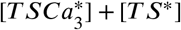 is approximated to be proportional to the value itself, *T*_*3*_ can be approximated to be linear with *F*_*b*_

*T*_4_ is a product of 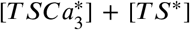 and the temporal differential of *h*_*p*_. As mentioned above, the power state 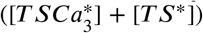 can be approximated to be linear with *F*_*b*_. The temporal differential of *h*_*p*_ can be deformed as follows.

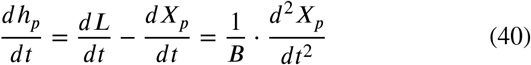

As explained in section 4.3.1, except for the initial ∽15 [ms] after the onset time of ejection, Eq. (40) reaches its steady

state. This means that 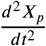 can be approximated as a small constant. As the value of is under 2% and increases to 8% of *F*_ext_ during the ejection phase, we approximate *F*_b_ as *F*_*ext*_ Afterward, by using *K*_1_ and *K*_2_ as constants, the following equation holds.

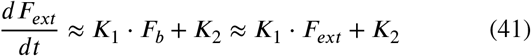

Afterward, by using Eqs. (3) and (2), the temporal differential of the half-sarcomere length 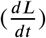 can be derived as follows.

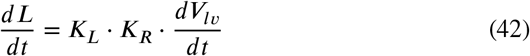

Here, the sum of the LV volume (*V*_*l,v*_) and the aortic volume (*V*_*a*_) during the ejection phase is denoted as the total volume (*V*_*tot*_).

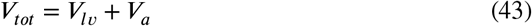

Subsequently,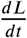 can be represented as follows.

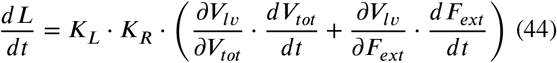

During the ejection phase, the temporal differential of the total volume 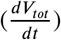 corresponds to the blood flow from the aorta to the peripshery, which can be represented as follows.

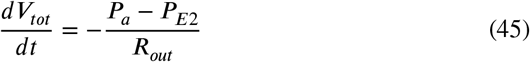

Moreover, as aortic resistance (*R*_*lo*_) is small, the following approximation holds.

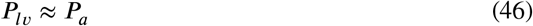

Now, 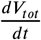, which is the first term on the right-hand side of Eq. (44), can be approximated as follows.

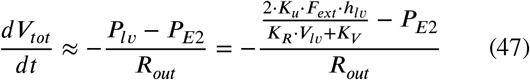

In addition, from Eqs. (1), (43), and (46), the total volume (*V*_*tot*_) can be approximated as follows.

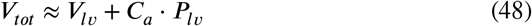

From Eq. (48), 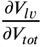 in the first term on the right-hand side of Eq. (44) can be obtained as follows.

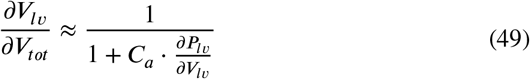

Similarly, by differentiating Eq. (48) with the LV wall tension (*F*_*ext*_), 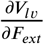 that appears in the second term on the right-hand side of Eq. (44) can be obtained as follows.

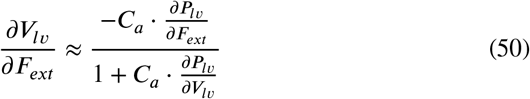

Here, the first term on the right-hand side of Eq. (44) is the product of Eq. (49) and Eq. (47). The changes in Eq. (49) with respect to changes in *F*_*ext*_ are small, while Eq. (47) consists of the component proportional to *F*_*ext*_ and constant. On the other hand, the second term on the right-hand side of Eq. (44) is the product of Eq. (50) and 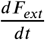 The changes in Eq. (50) with respect to changes in *F*_*ext*_ are small. Thus, this term can be approximated as proportional to 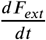, as changes in other terms are small with respect to changes in *F*_*ext*_ and *L*.

In summary, the following approximation holds with the constants *K*_3_, *K*_4_, and *K*_5_.

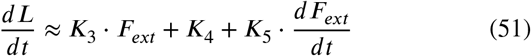

By using this equation and Eq. (41), the following equation holds.

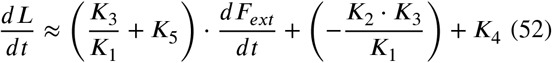

Therefore, except for the initial transient period, the relation between 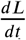 and 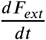 approximately satisfies Eq. (32) during the ejection phase. This occurs under the assumption that the relation between *L* and *F*_*ext*_ satisfies Eq. (29).

In the simulation results at 50 and 100 [ms] after the onset time of ejection, we can find a highly linear relation between 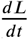 and 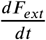, as shown in Figure 8. Similarly, at 280 [ms] after the onset time of ejection, we also can find a highly linear relation as shown in Figure 9.

**Figure 8:**
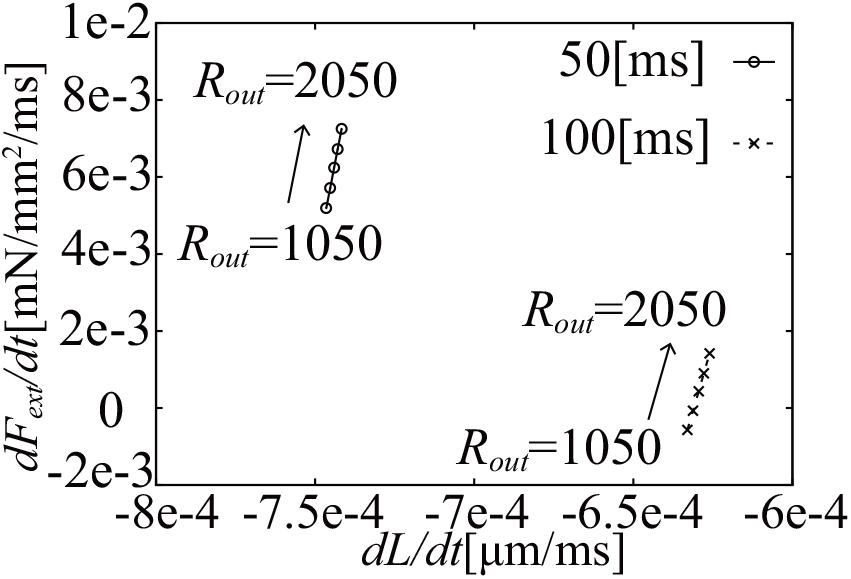
The relation between 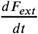 and 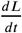 at 50, 100 [ms] after the onset time of ejection

**Figure 9:**
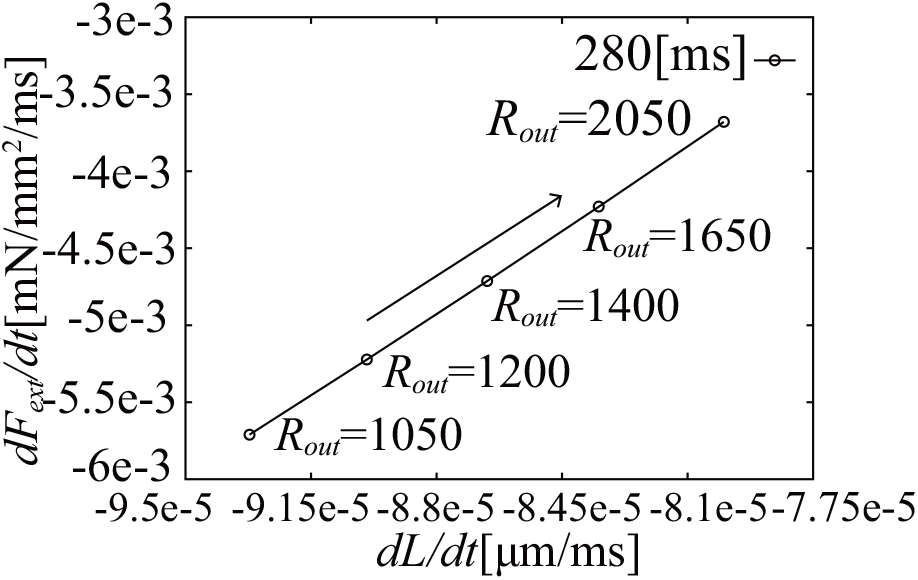
The relation between 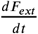 and 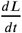 at 280 [ms] after the onset time of ejection

Thus, from the above analysis, we can understand why the isochronous LV wall tension (*F*_*ext*_) -half-sarcomere length (*L*) relation remains linear under the variation of the afterload (*R*_*out*_) during the ejection phase; that is, the characteristics come from the combination of the characteristics of the cardiac muscle and compliance of the LV and aorta.

### 4.4 Elastance of the time-varying elastance model (TVEM)

The Time Varying Elastance Model (TVEM) is one of the major simple LV model often used in hemodynamic models. In the model, a time-dependent elastic function (*E*(*t*)) and the constant zero-pressure fill volume *V*_0_ are used to relate LV pressure and LV volume as follows.

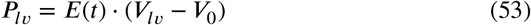

Thus, in the TVEM, *E*_*inst*_ and *E*_*load*_ always acquire the same value of *E*(*t*).

## 5. Discussion

### 5.1 Relation between *E*_*max*_ and *E*_*load*_

*E*_max_ is the slope of ESPVR proposed as an index of carcontractility [43, 41]. From the simulation results, the alue of *E*_*load*_ at 280 [ms] after the onset time of the ejection, which was close to that of the end-systole, was close to *E*_*max*_. With our hemodynamic model, the end-systolic time slightly changed with respect to the afterload (*R*_*out*_); thus, the isochronous *F*_*ext*_ -*L* relation at a fixed time close to the end-systole time became slightly different from the ESPVR. However, from the simulation results, the end-systolic time was almost linear with the LV end-systolic pressure (*P*_*l,v*_); thus, the end-systolic points were in a line and the difference between the line and the isochrones of *F*_*ext*_ -*L* was very small.

### 5.2 Relationship between Force-Velocity Relation and *E*_*load*_

In our analysis, the initial isochronous *P*_*l,v*_ -*V*_*l,v*_ relation slope was determined by the initial transient changes in the half-sarcomere length and cellular contraction force, as explained in section 4.3.1. Moreover, the slope was approximately maintained during the ejection phase. In this period, the sarcomere shortening velocity was determined by the mechanical load to the cell corresponding to the force-velocity relation (FVR); thus, the slope, i.e., *E*_*load*_ was strongly related to the FVR characteristics but not to the cardiac tissue contractility.

Since the shape of FVR is known to be physiologically non-linear [10, 14, 40], the isochronous PV relation should become non-linear. However, as the range of the force variation is quite small, FVR becomes approximately linear, resulting in a linear isochronous PV relation.

On the other hand, the positive inotropic effect, such as the noradrenergic stimulation, increases *E*_*max*_, which corresponds to the increase in *E*_*load*_ . This can be explained by the changes in FVR under the positive inotropic effect. As reported in Sonnenblick et al. [10, 14, 40], the positive inotropic effect changes the properties of FVR. In our model, we did not include the inotropic effect. Thus, this aspect must be considered in the future model.

### 5.3. Time course of *E*_*inst*_ and *E*_*load*_

From Table 7, we can observe that *E*_*inst*_ slightly increases and then decreases, while *E*_*load*_ monotonically decreases during the ejection phase. As shown in section 4.2, *E*_*inst*_ directly reflects cellular contractility. Thus, the time course of *E*_*inst*_ is similar to the cellular contraction force (*F*_*b*_). Additionally, as shown in section 4.3, *E*_*load*_ reflects the characteristics of FVR. Therefore, the time course of *E*_*load*_ is considered to reflect the time course of the ratio between 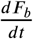 and 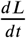

### 5.4 Comparison between the simplified hemodynamic model and TVEM

Using a model in which the TVEM is used instead of the simplified hemodynamic model of the LV compartment, which was used by Heldt et al. [18], we evaluated the PV loops under the following peripheral resistance conditions:

*R*_*out*_ = 1050, 1200, 1400, 1650, and 2050 [mmHg *⋅* ms/mL]. The resulting PV loops and isochronous of the *P*_*l,v*_ -*V*_*l,v*_ relations at 5, 50, 100, 150, and 200[ms] after the onset time of ejection are shown in Figure 10. Moreover, the time course of elastance (*E*) is shown in Figure 11. As shown in Figure 10, the slope of the isochronous *P*_*l,v*_ -*V*_*l,v*_ relation monotonically increases. In addition, as shown in Figure 11, *E* also increases monotonically. This is determined by the elastance function used for TVEM. As explained in section 3.2, the time course of the *E*_*load*_ of our simplified hemodynamic model slightly decreases with time, which is different from that of the TVEM.

**Figure 10:**
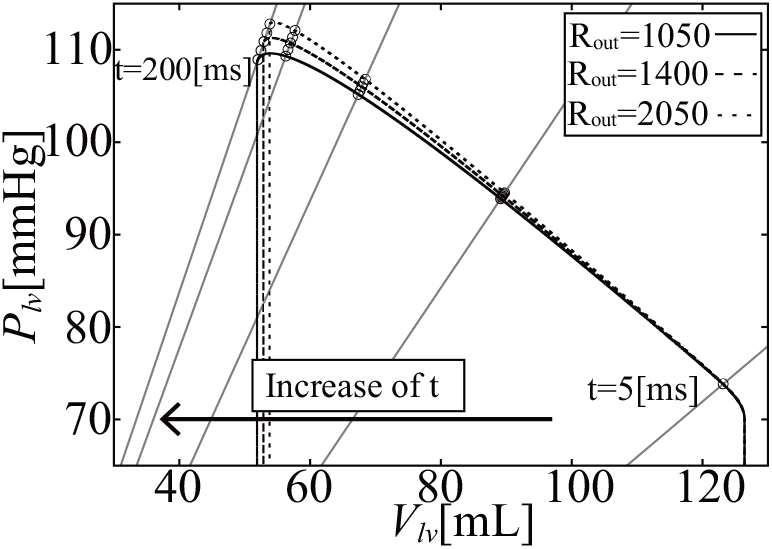
Isochronous *P*_*l,v*_ -*V*_*l,v*_ relations during the ejection phase when using TVEM

**Figure 11:**
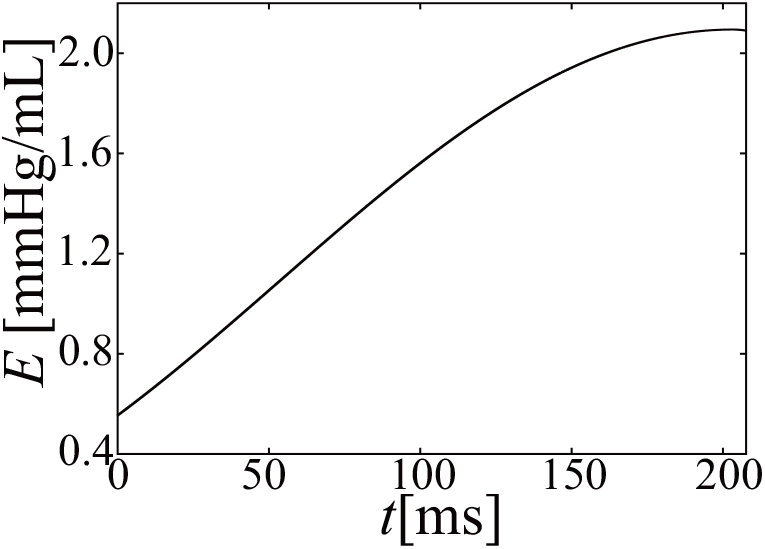
Time course of elastance (E) after the onset time of ejection when using TVEM

For the animal experiment, Nishioka et al. [29] measured the isochronous PV relation in dogs and the results came close to our simulation model. That is, the slope of the isochronous PV relation did not change with time, but the volume intercept changed. However, in their experiment, pressure and volume at the onset time of ejection were not fixed. Thus, we cannot say that their results correspond to those of our simulation, but there are some similarities with our results.

### 5.5 Influence of simplification of the hemodynamic model

To simplify the mathematical analysis, we simplified the hemodynamic model proposed by Utaki et al. [49, 45] for two points. First, LV wall thickness was fixed at a constant length, and second, the relation between the LV internal radius (*R*_*l,v*_) and LV volume (*V*_*l,v*_) was simplified to linear. However, these simplifications may reduce the accuracy of the model.

If we use the original *h*_*l,v*_, which is proportional to the cellular contraction force (*F*_*b*_), the *E*_*inst*_ equation (Eq. (20)) becomes as follows.

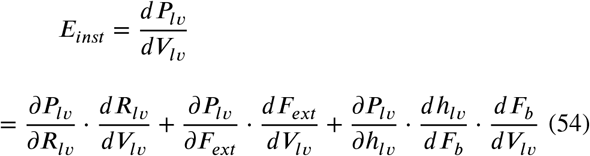

Comparing Eq. (20) with Eq. (54), we can see that the third term on the right-hand side of Eq. (54) is ignored in our analysis. This term may lower the accuracy of the simulation results; however, from the simulation results shown in Figure 2, the simplified hemodynamic model still can reproduce physiological hemodynamics.

Additionally, as shown in Eq. (42), the relation between the half-sarcomere length (*L*) and the LV volume (*V*_*l,v*_) is simplified to a linear relation in our model, while the relation becomes non-linear in dogs, as reported by Rodriguez et al. [31]. In this case, the isochronous PV relation may become non-linear. In the case of rats and sheep, the isochronous PV relation is reported as non-linear [11, 24]. Thus, part of the difference with our simulation results may be caused by this simplification.

### 5.6 Limitations of the analysis

In the analysis of step 2 (section 4.3.2), the slope of the relation between the LV wall tension (*F*_*ext*_) and half-sarcomere length (*L*) is assumed to be fixed, while as shown in the simulation results (Table 7), the slope slightly decreases during the ejection phase. We could not derive an analytical solution without a fixed slope value; thus, in this aspect, we need further analysis of the mathematical equations of the model.

In the calculation of 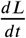 from Eq. (44), 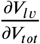 and 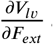 are calculated from Eqs. (49) and (50). Note that, these equations cannot be calculated when 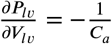.

## 6. Conclusions

In this paper, we evaluated the simulation results of the proposed hemodynamic model that included a detailed cellular contraction model. By defining two different LV elastances: 1) instantaneous elastance and 2) load-dependent elastance, we were able to evaluate these elastances from the simulation model and found that these elastances showed markedly different time courses. That is, the instantaneous elastances showed a bell-shaped curve corresponding to the cellular contraction force, while the load-dependent elastance hardly changed with time. We then analyzed the mechanism that determines these elastances from the model equations, and found that the instantaneous elastance directly coincided with the cellular contraction force, while the load-dependent elastance was determined by the characteristics of both the instantaneous isovelocity-shortening and the force-velocity relation of cardiac cells. The slope of the isochronous pressure-volume relation was mainly determined by the velocity-dependent force drop characteristics in the instantaneous shortening. Moreover, the linear relation between the isochronous pressure and volume was based on the characteristics that the relation between the temporal differential of the cellular contraction force and the cellular shortening velocity becomes linear, which comes from the combined characteristics of the LV and aortic compliances.

## Supporting information

Program source code

