## Supplementary figures and images for "Mathematical Analysis of Left Ventricular Elastance with respect to Afterload Change During Ejection Phase"

### 0-2-01_threeComponents.png

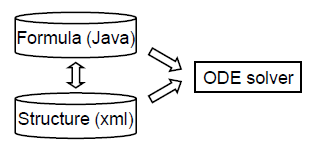

### 0-2-02_Java.png

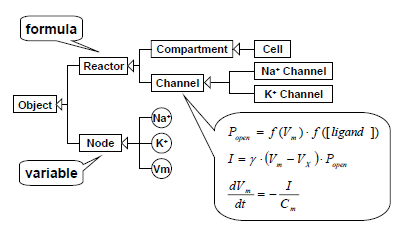

### 0-2-03_Xml.png

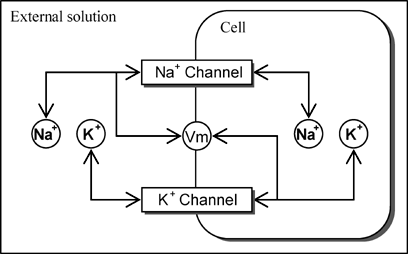

### 1-1-1_WorkspaceLauncher.png

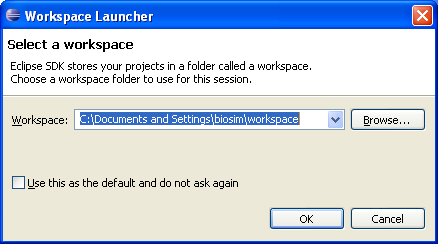

### 1-1-2_WelcomeToEclipse.png

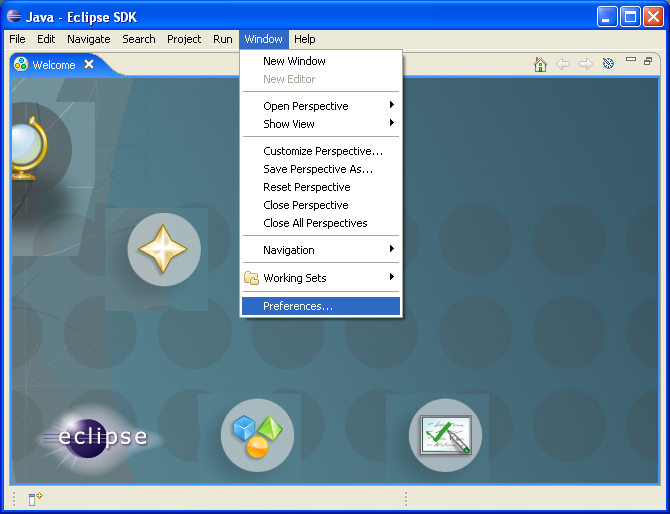

### 1-1-3_setUTF-8.png

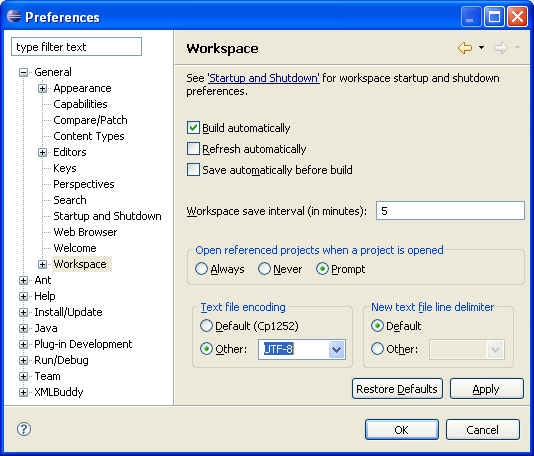

### 1-1-4.png

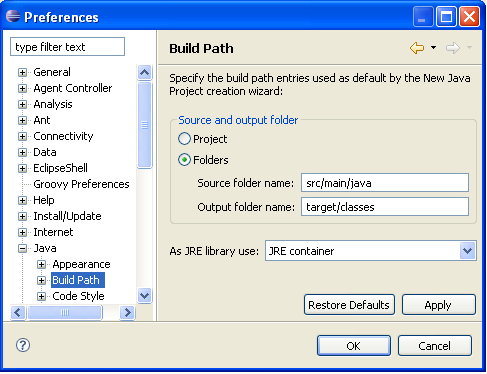

### 1-1-5.png

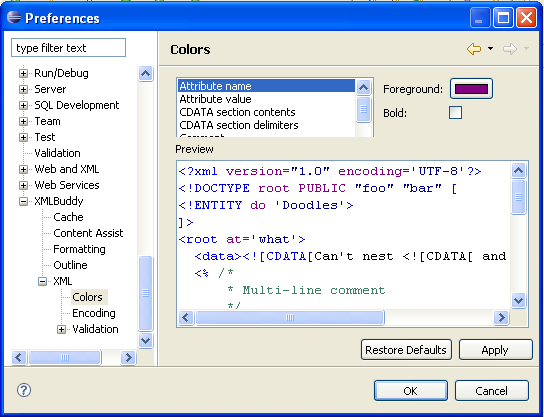

### 1-1-5_UserLibraries.png

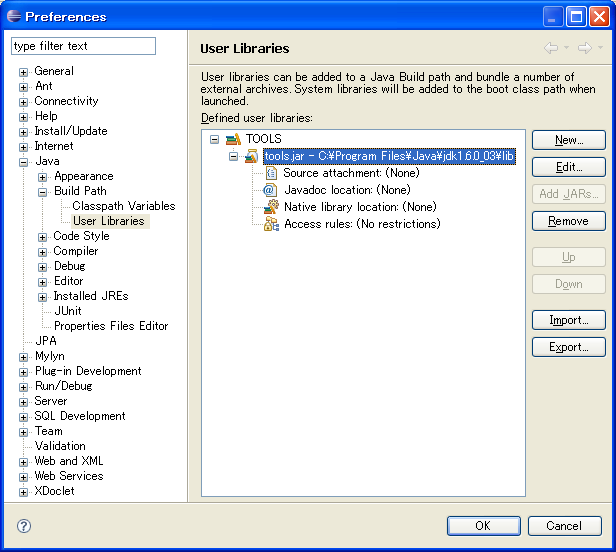

### 1-2-1.png

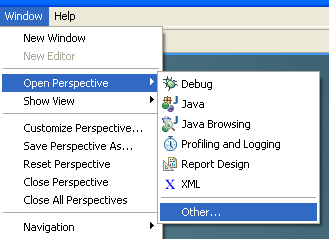

### 1-2-2.png

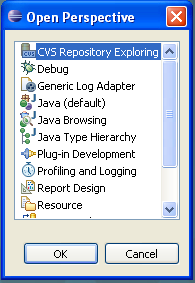

### 1-2-3.png

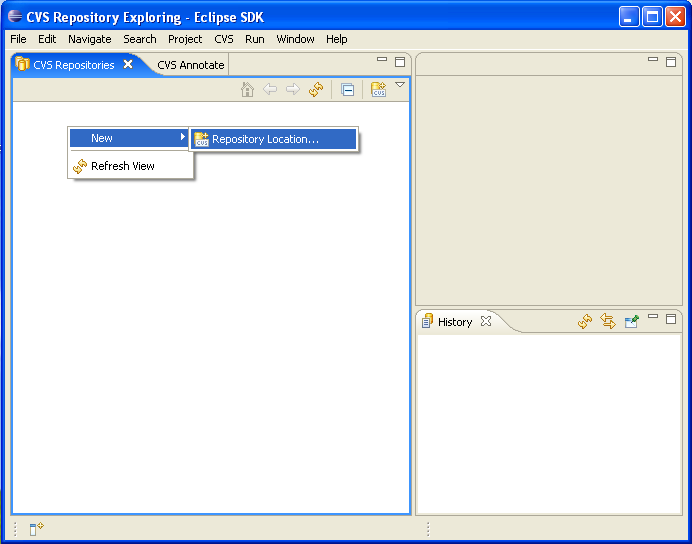

### 1-2-4.png

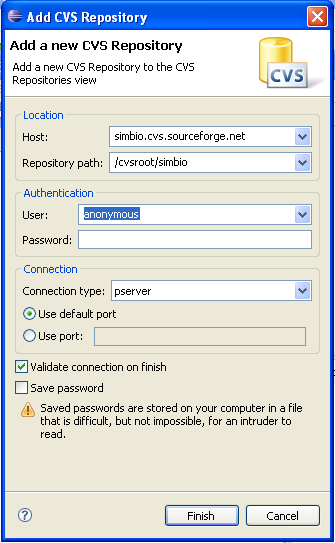

### 1-2-5.png

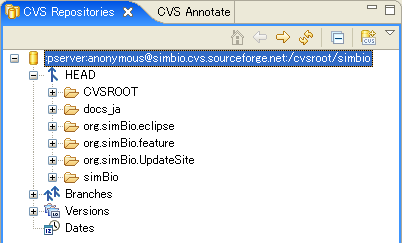

### 1-2-6.png

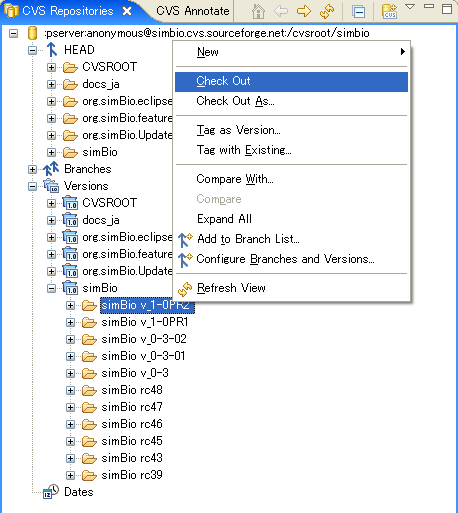

### 1-2-7.png

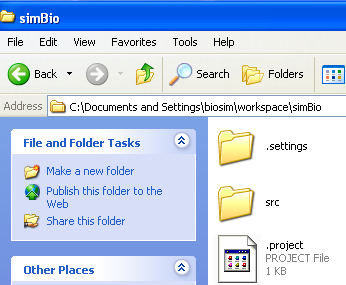

### 1-2-8.png

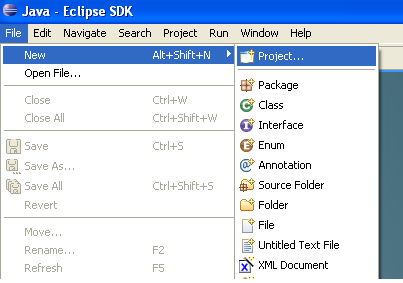

### 1-2-9.png

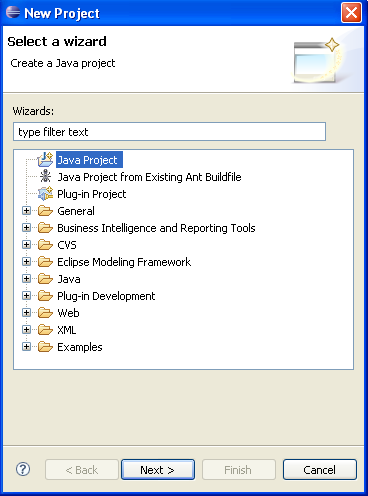

### 1-2-11.png

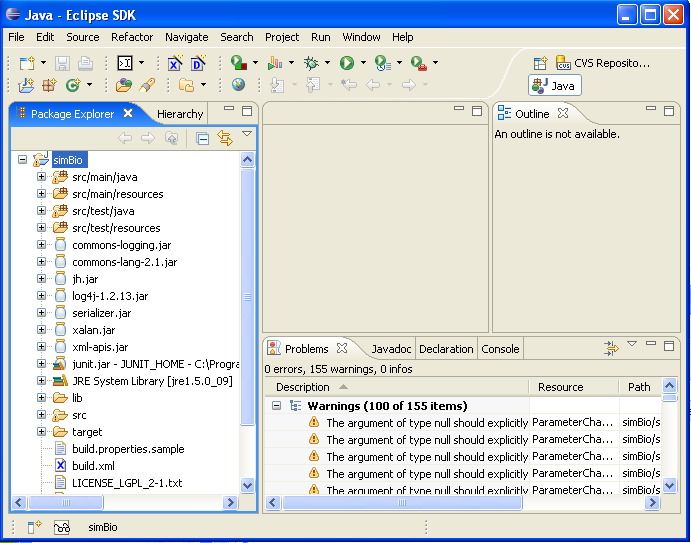

### 1-2-12.png

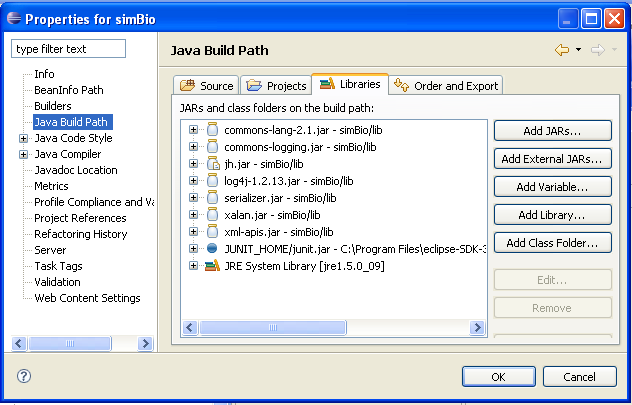

### 1-2-13_JDKComplianceSettings.png

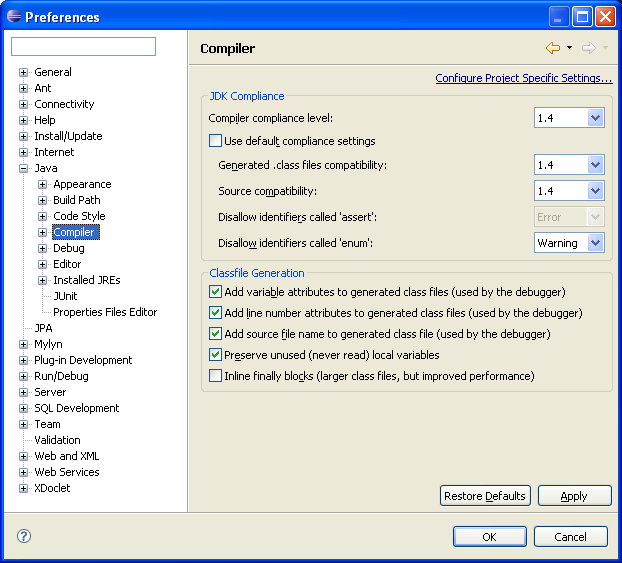

### 1-2_CVS_proxy.png

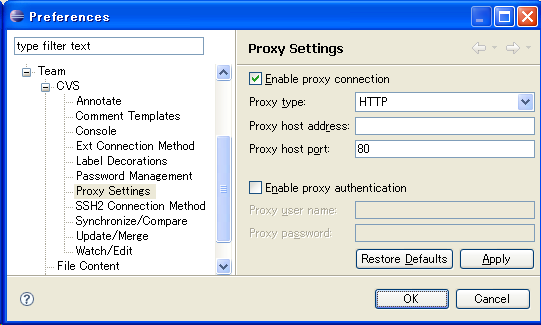

### 1-2_NewJavaProject.png

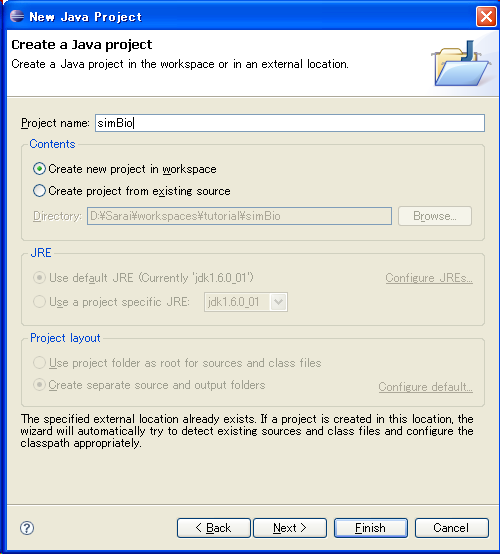

### 1-3-1.png

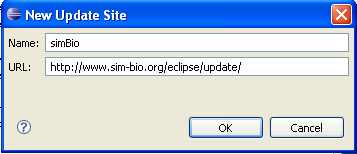

### 1-3-2.png

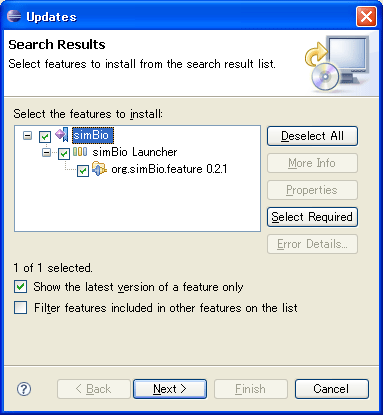

### 1-3-3.png

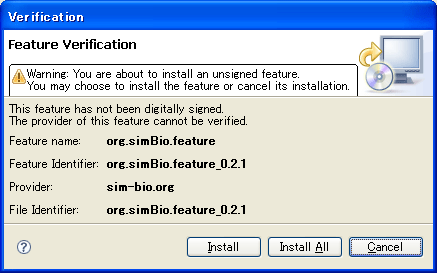

### 1-3_License.png

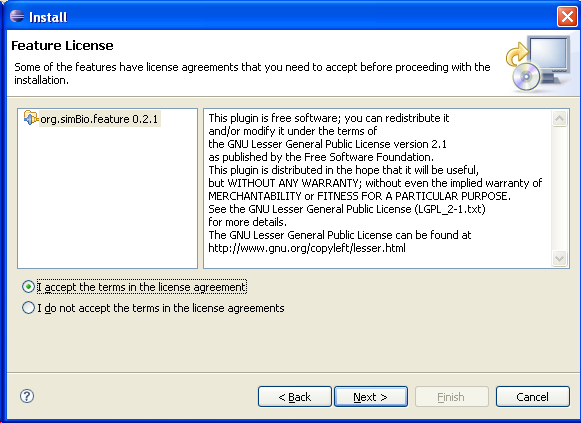

### 1-3_UpdateSiteToVisit.png

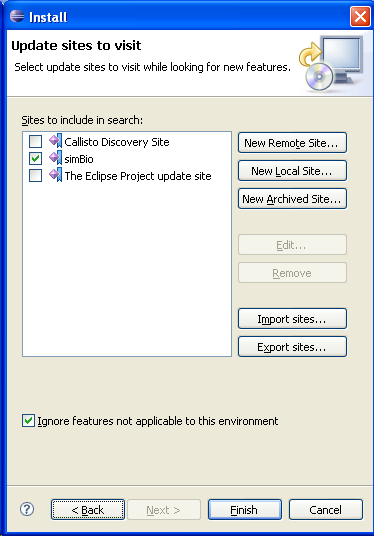

### 2-1-01_RunOnGUI.png

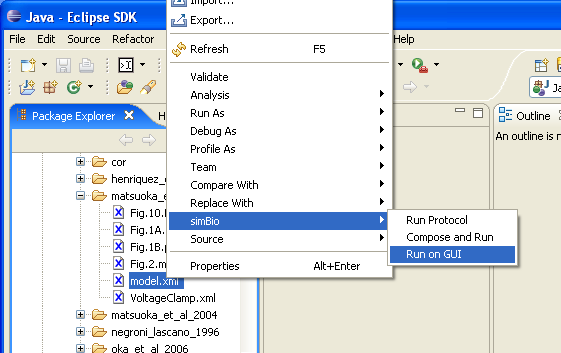

### 2-1-02_matsuoka_et_al_2003.png

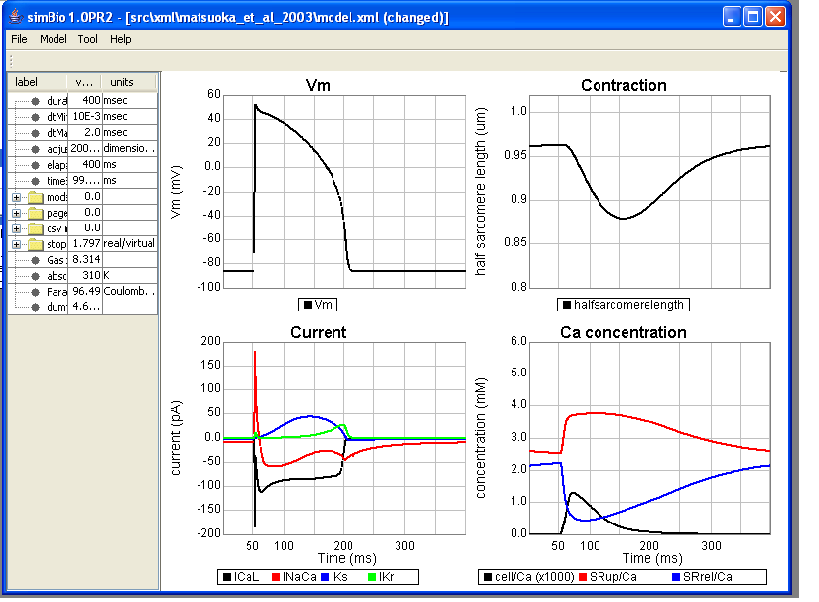

### 2-1-03_Coolbar.png

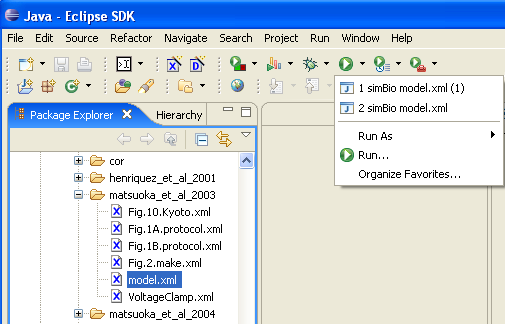

### 2-1-04_Menu.png

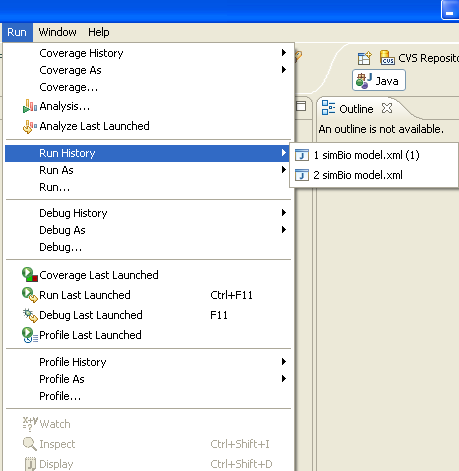

### 2-1-05_editToolbar.png

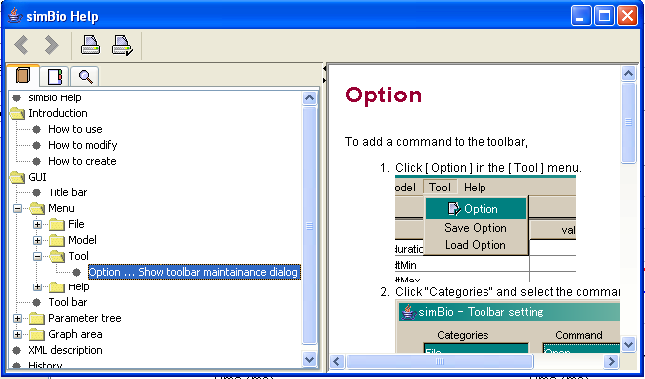

### 2-1-06_myocyte_mini.png

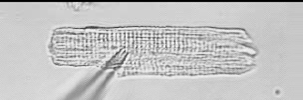

### 2-1-07_interaction.png

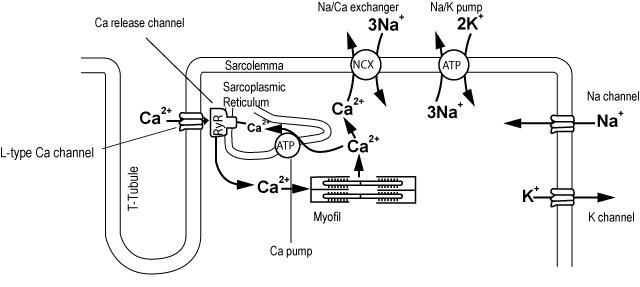

### 2-1-08_NoAP.png

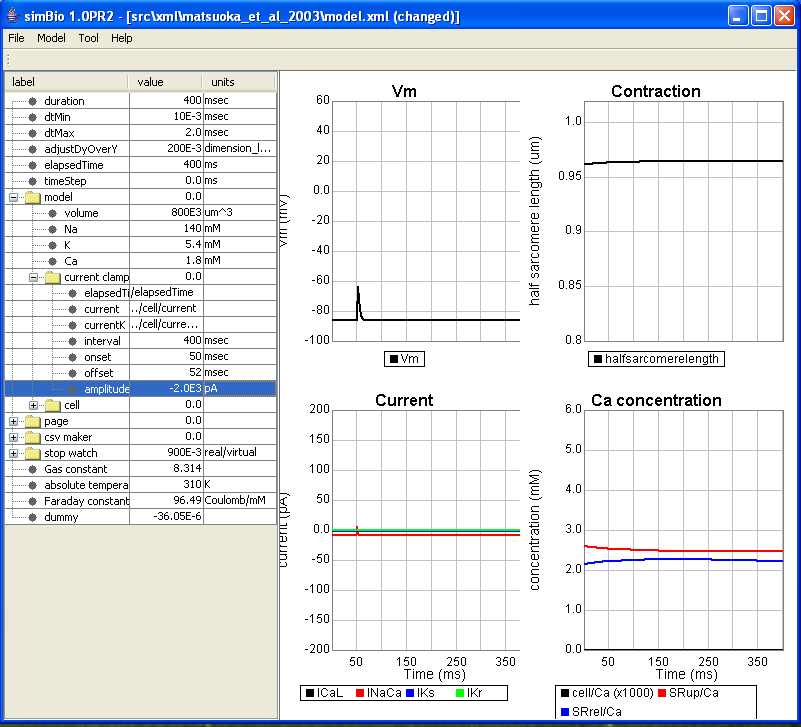

### 2-1-09_ActionPotentials.png

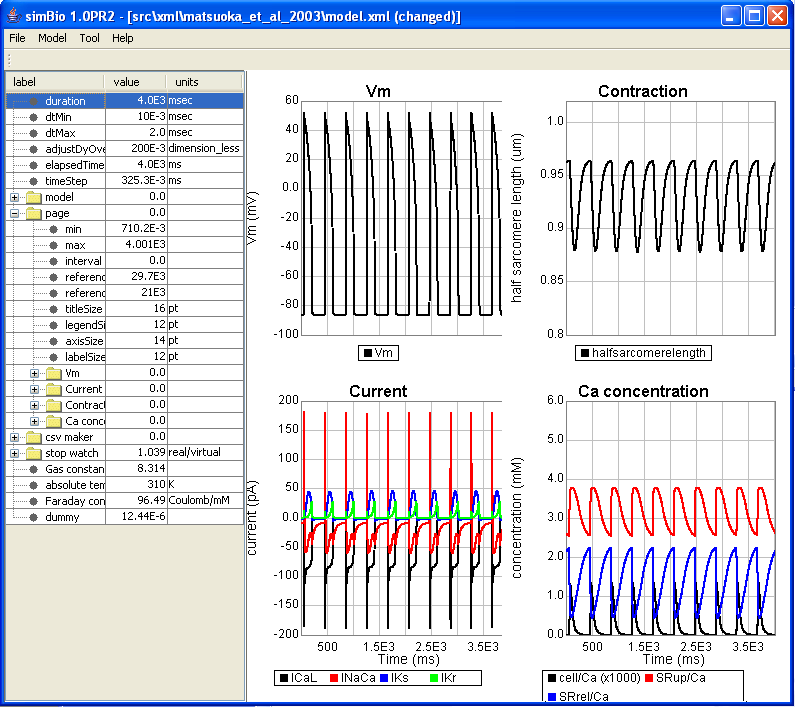

### 2-1-10_AllOrNone.png

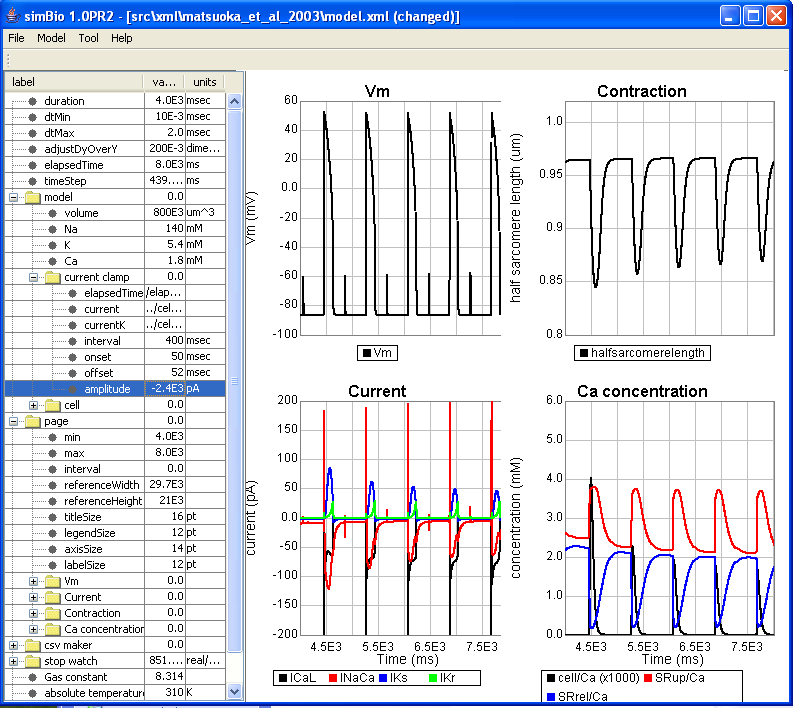

### 2-1-11_PositiveStairCase.png

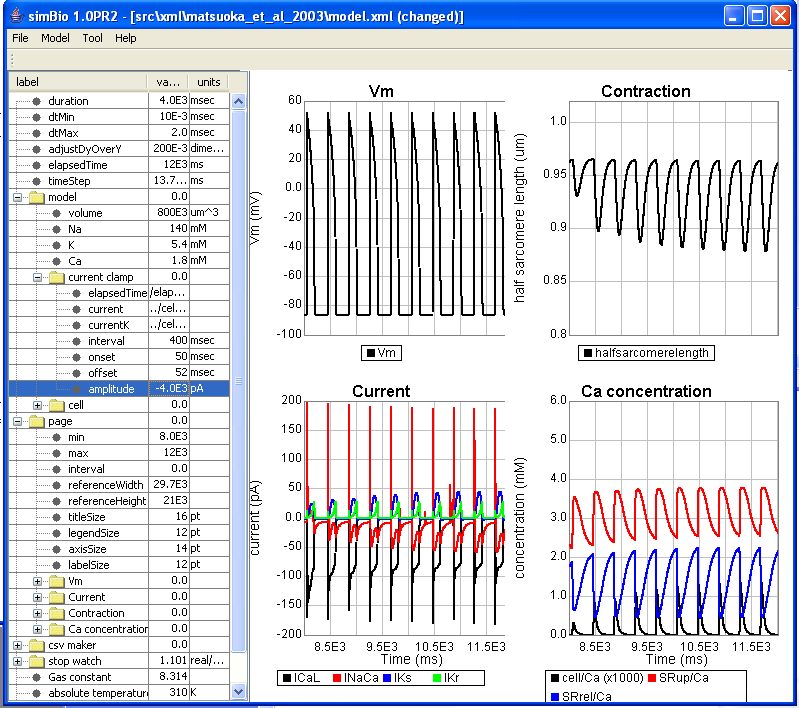

### 2-1-12_LowKo.png

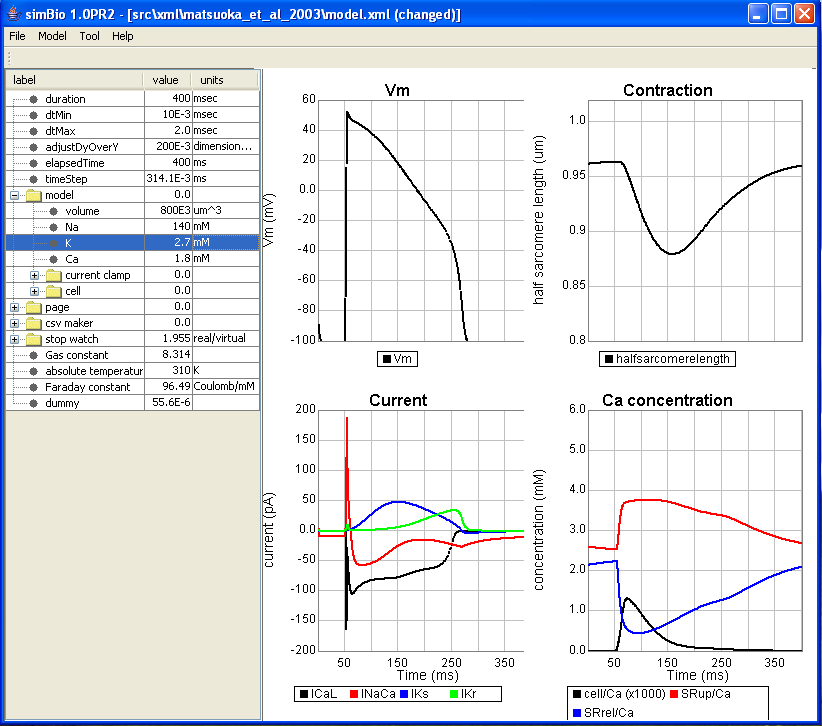

### 2-1-13_sarai_et_al_2003.png

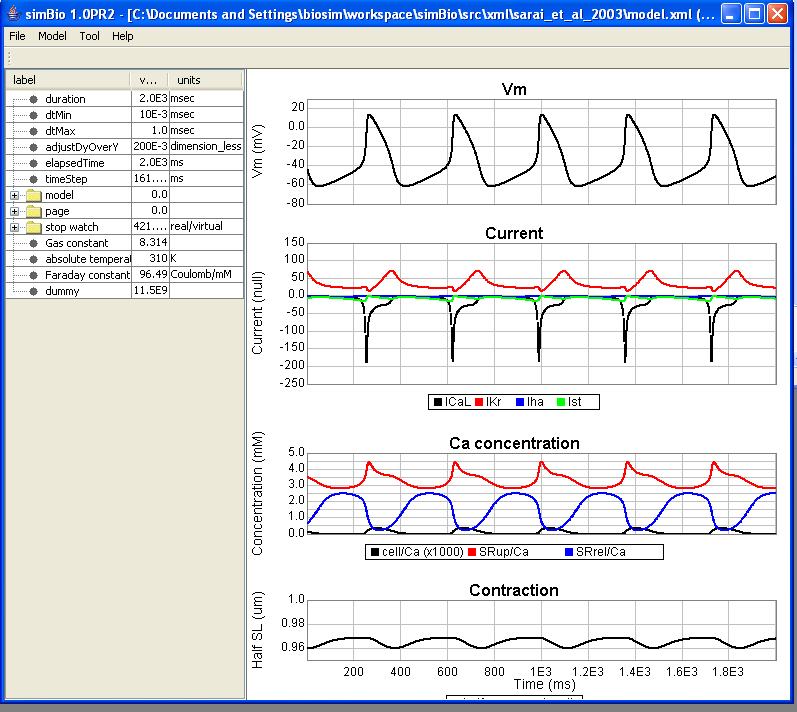

### 2-1_parameter.png

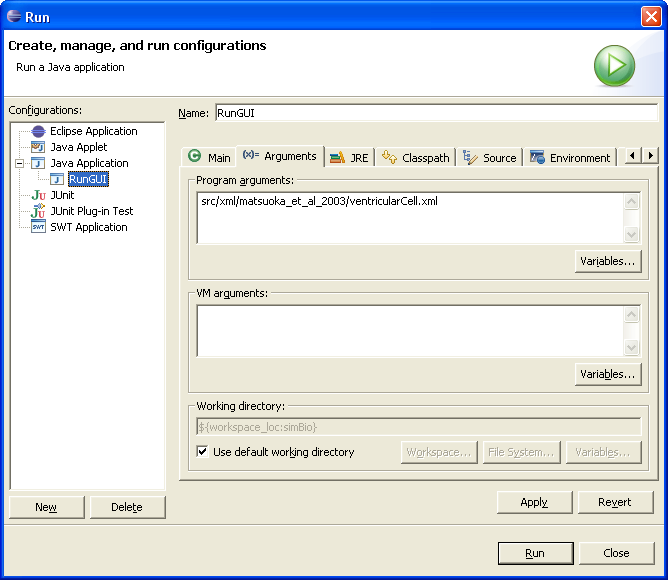

### 2-1_Run.png

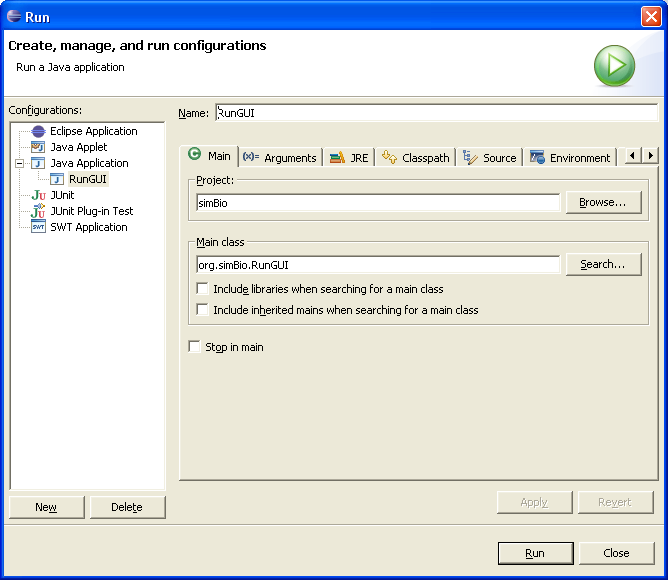

### 2-1_RunGUI.png

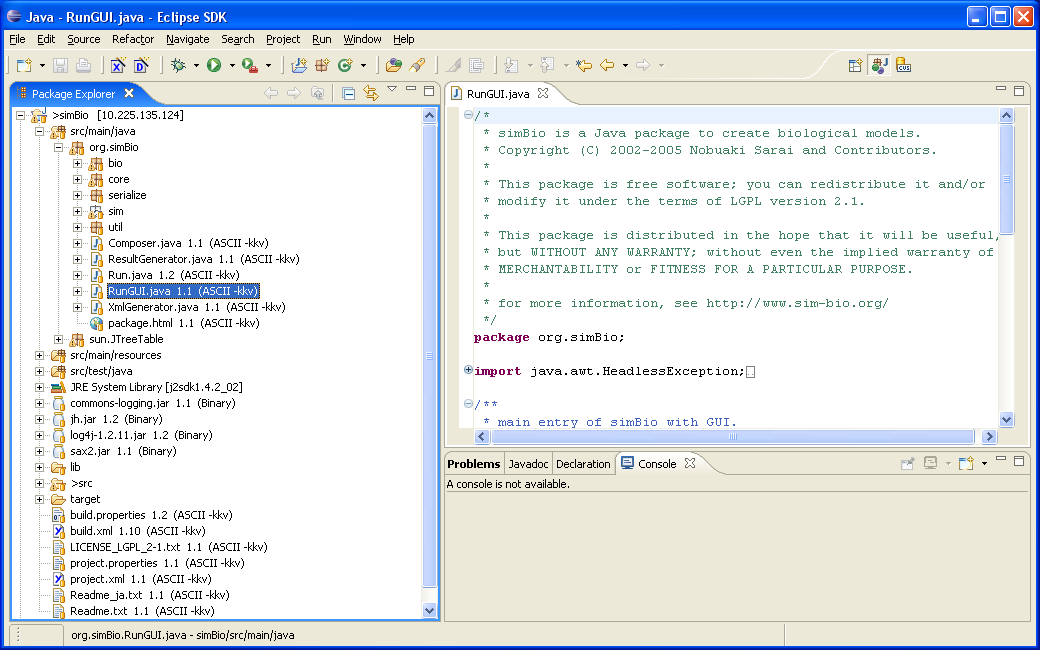

### 2-2-1_basicXML.png

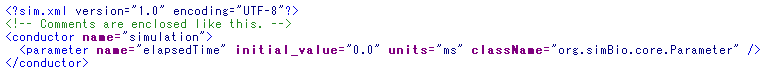

### 2-2-2_xmlTree.png

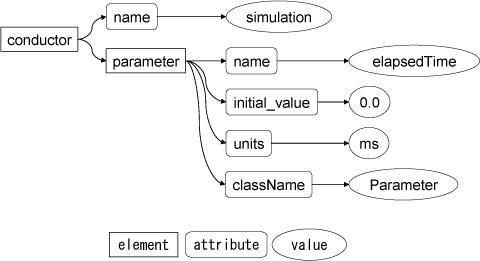

### 2-2-3_instanceTree.png

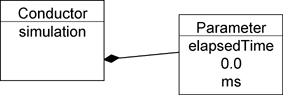

### 2-2-4_tutorial_property.png

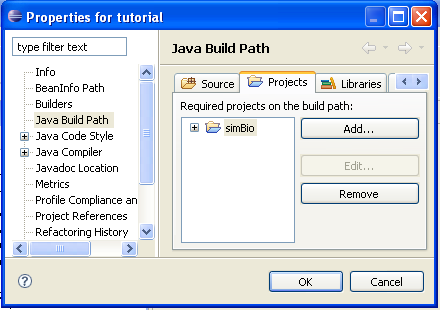

### 2-2-5_tutorial_project.png

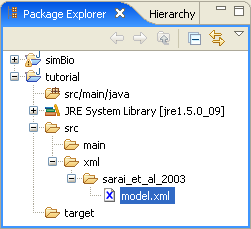

### 2-2-6_edit_modelK.png

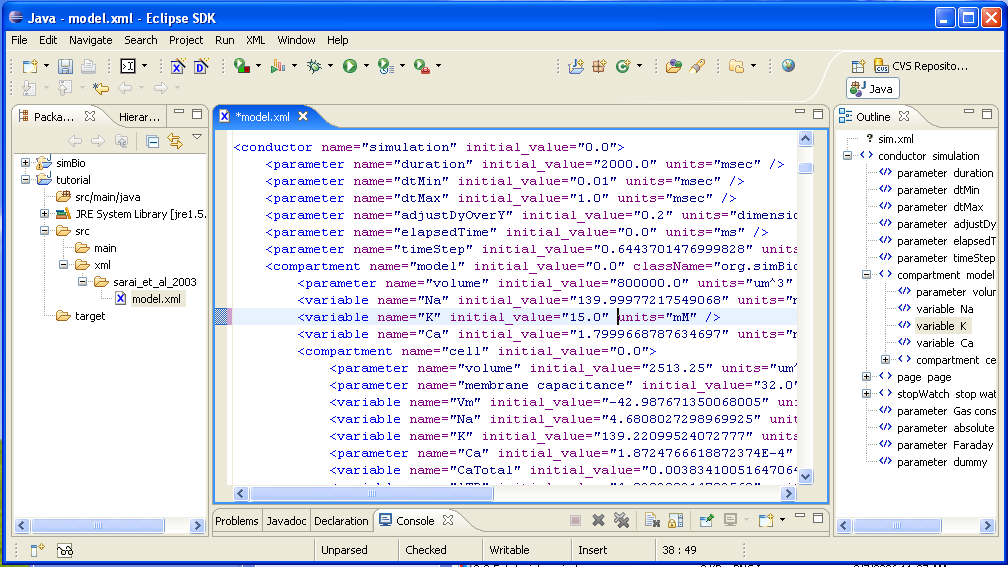

### 2-2-7_modelK15.png

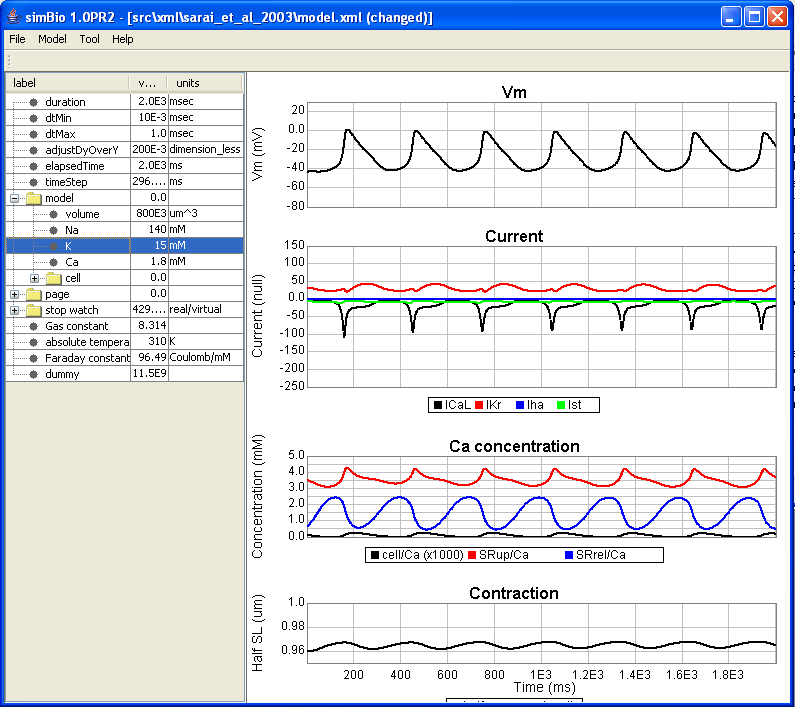

### 2-2-8_modelK5.4.png

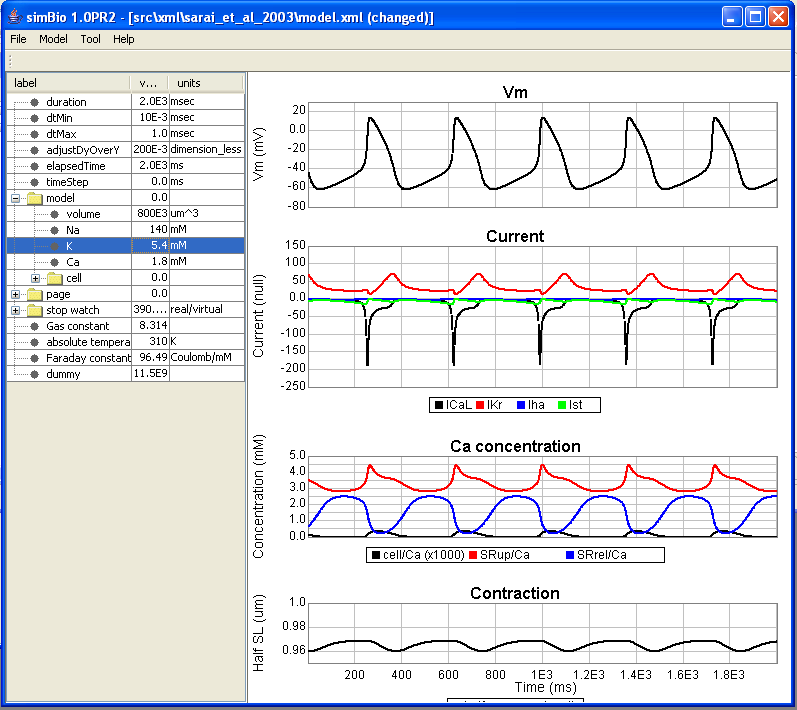

### 2-3-1_replace.png

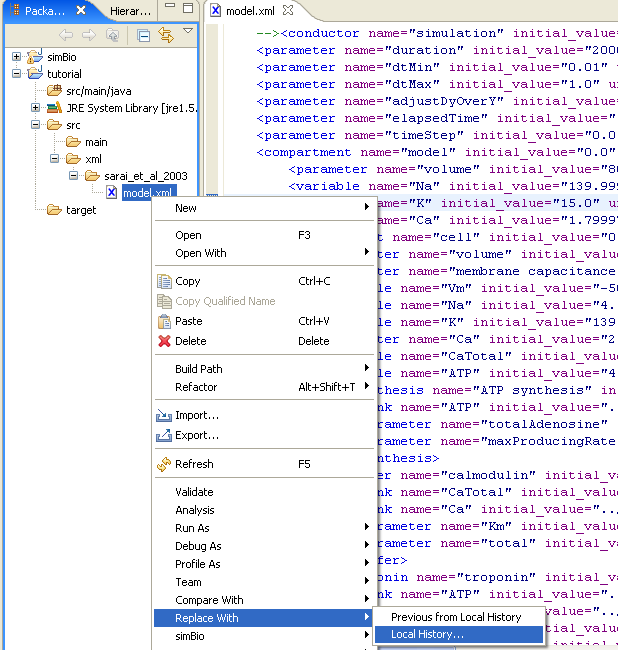

### 2-3-2_deleteINaK.png

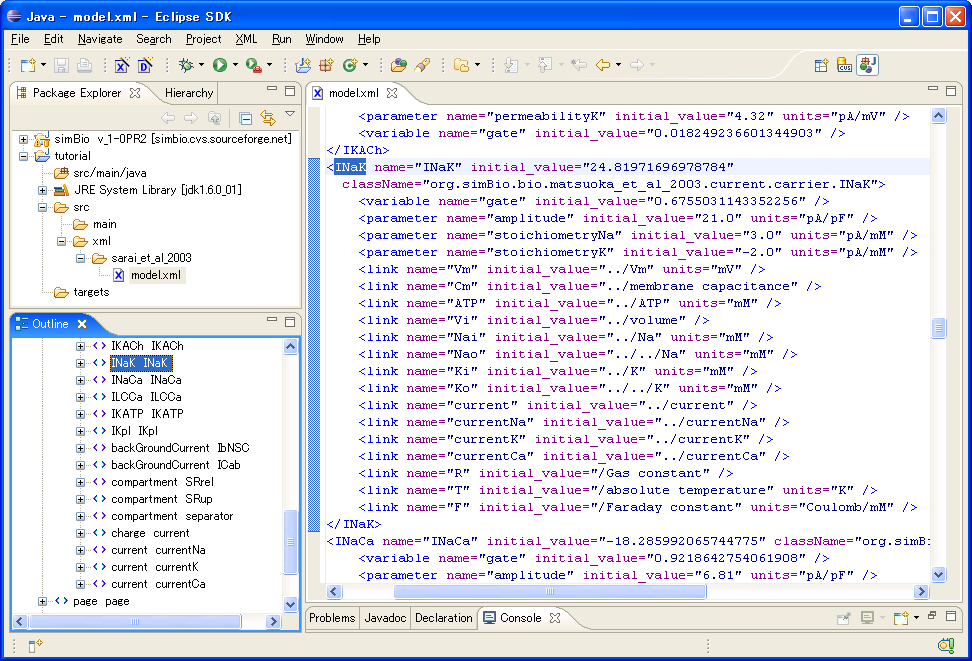

### 2-3-3_NoINaK.png

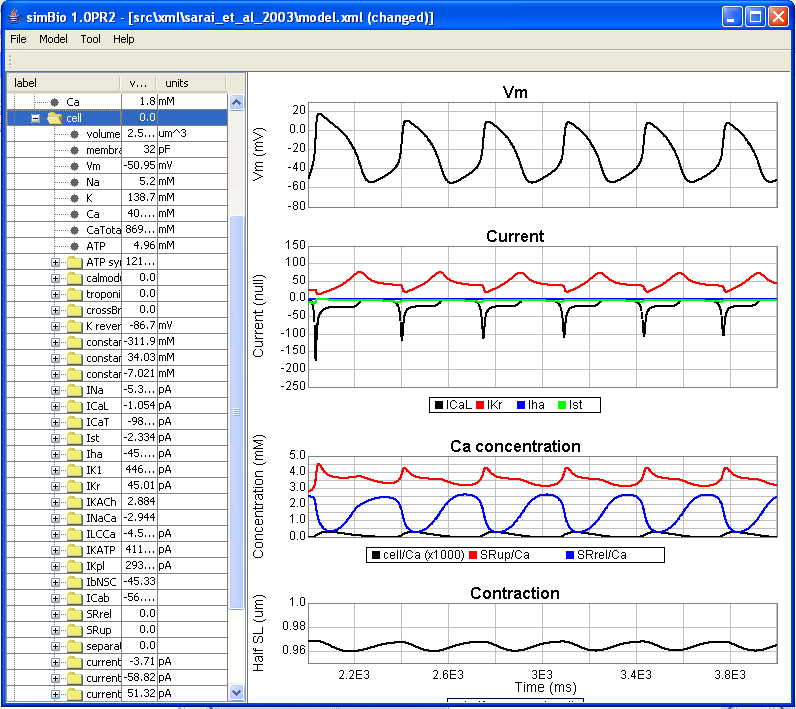

### 2-3-4_Control.png

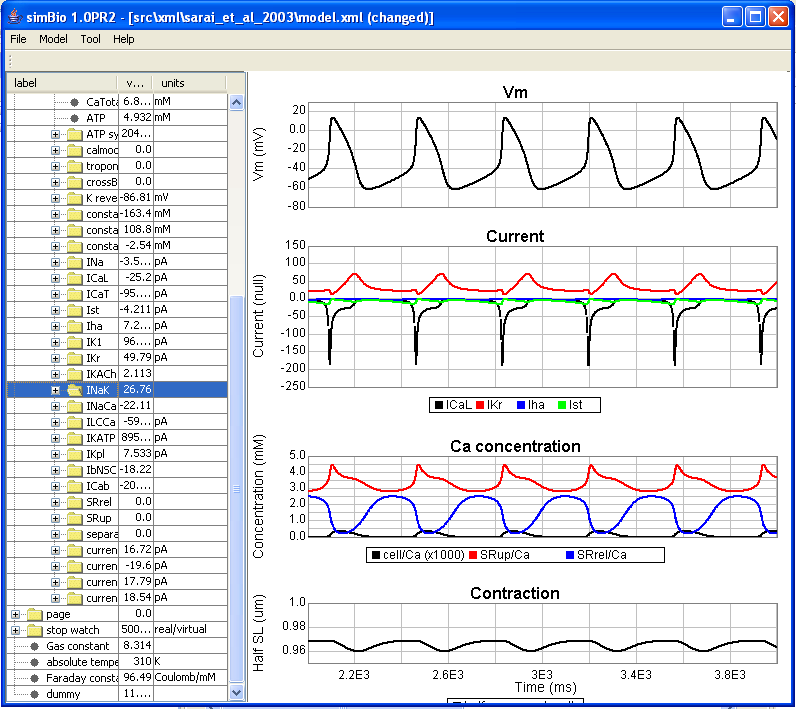

### 2-4-1_help.png

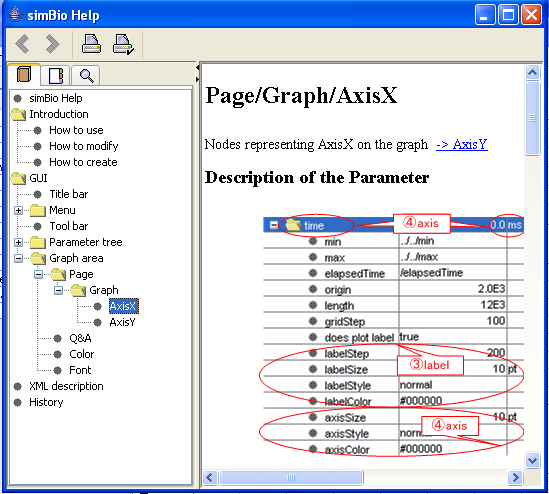

### 2-4-2_explanation.png

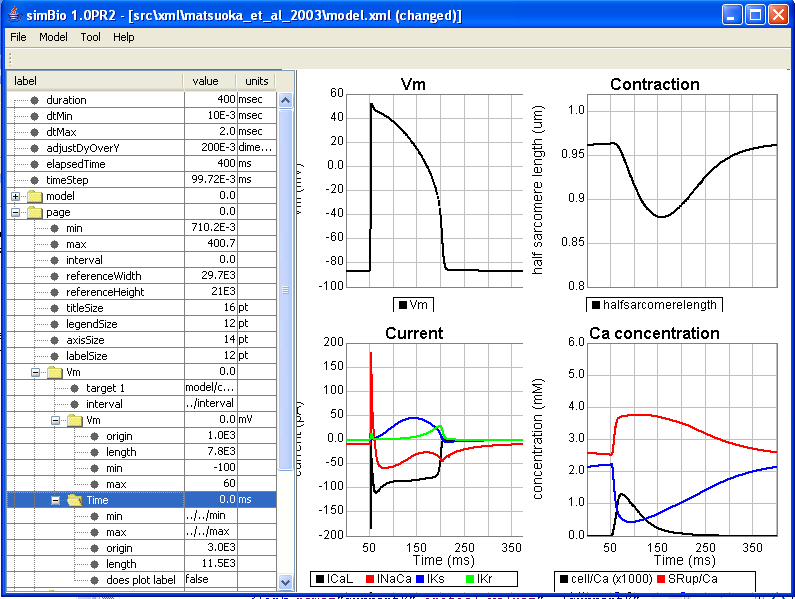

### 2-4-3_page.png

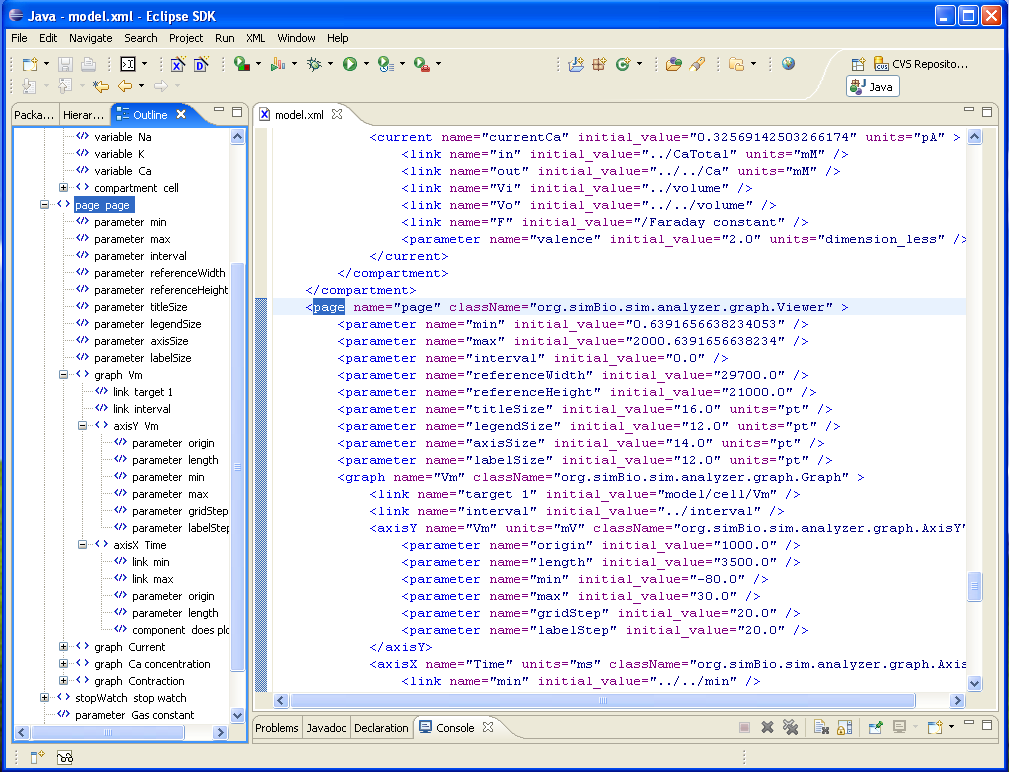

### 2Dcontour.png

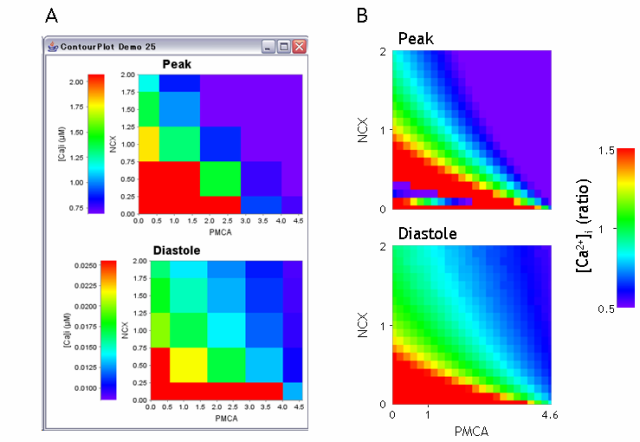

### 3-1-01.png

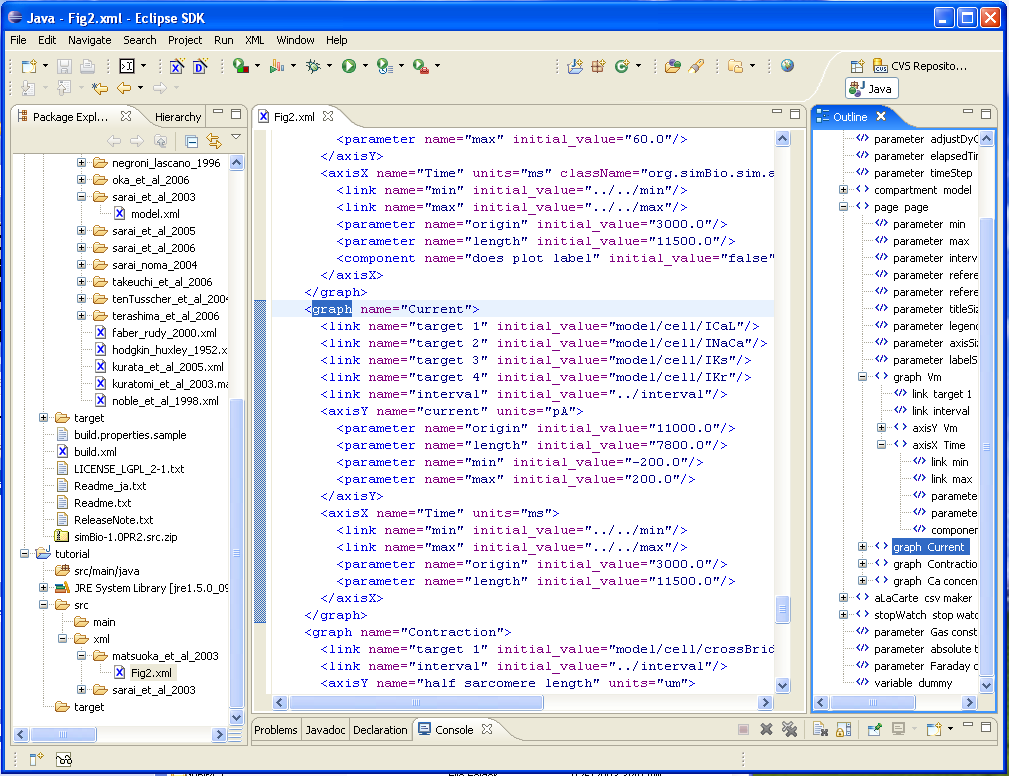

### 3-1-02.png

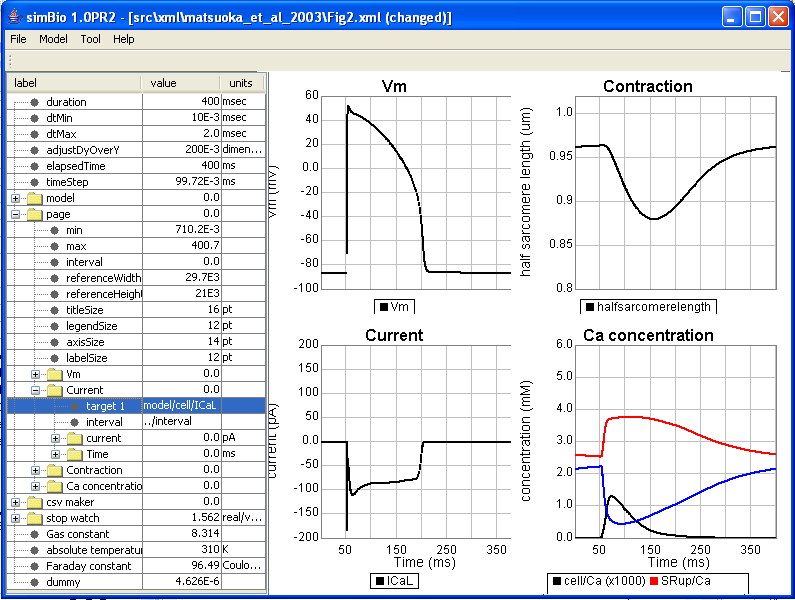

### 3-1-03.png

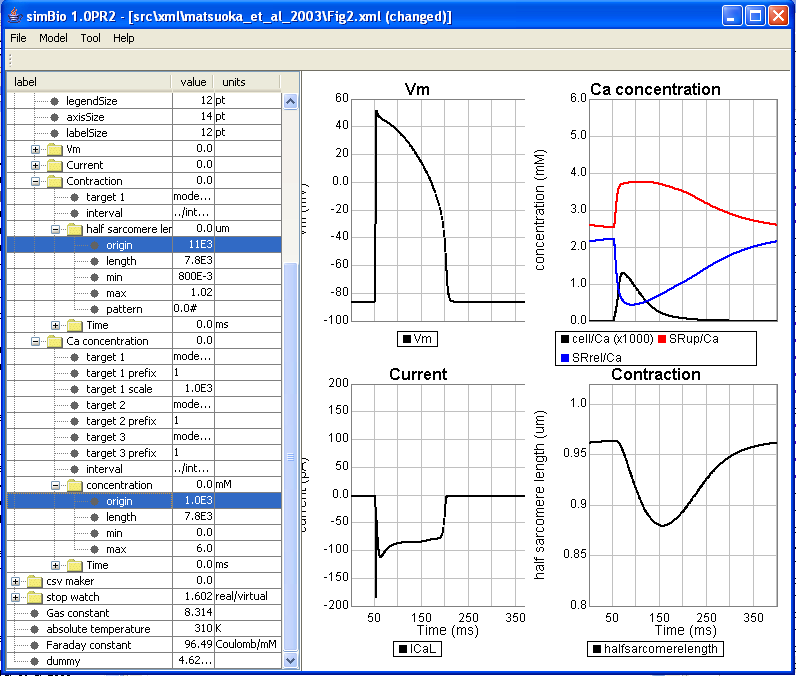

### 3-1-04_NameConflict.png

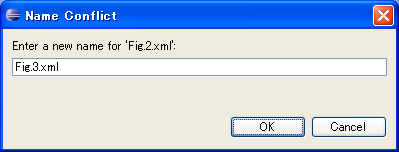

### 3-1-05.png

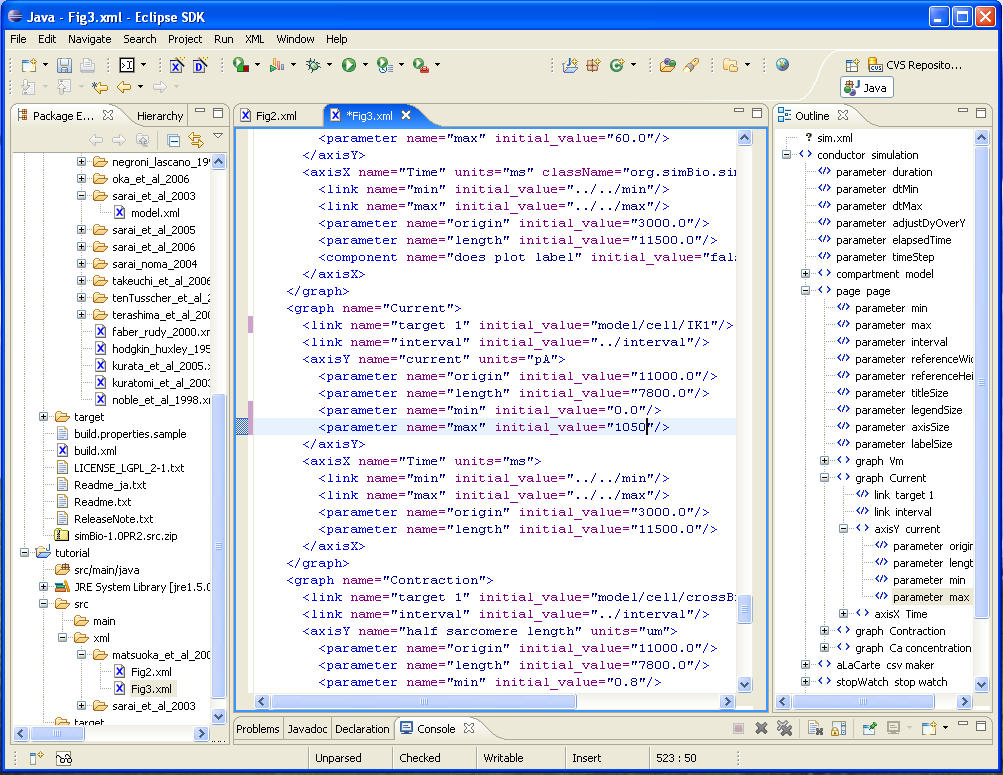

### 3-1-06.png

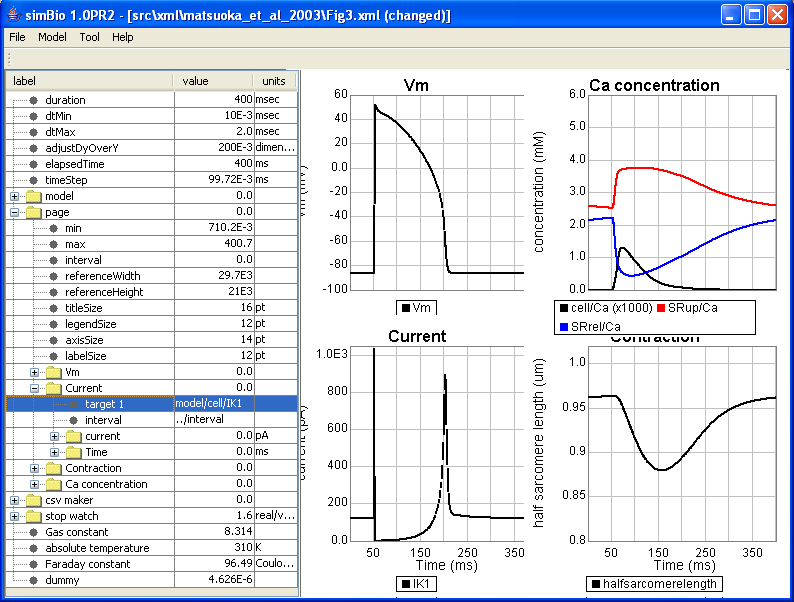

### 3-1-07.png

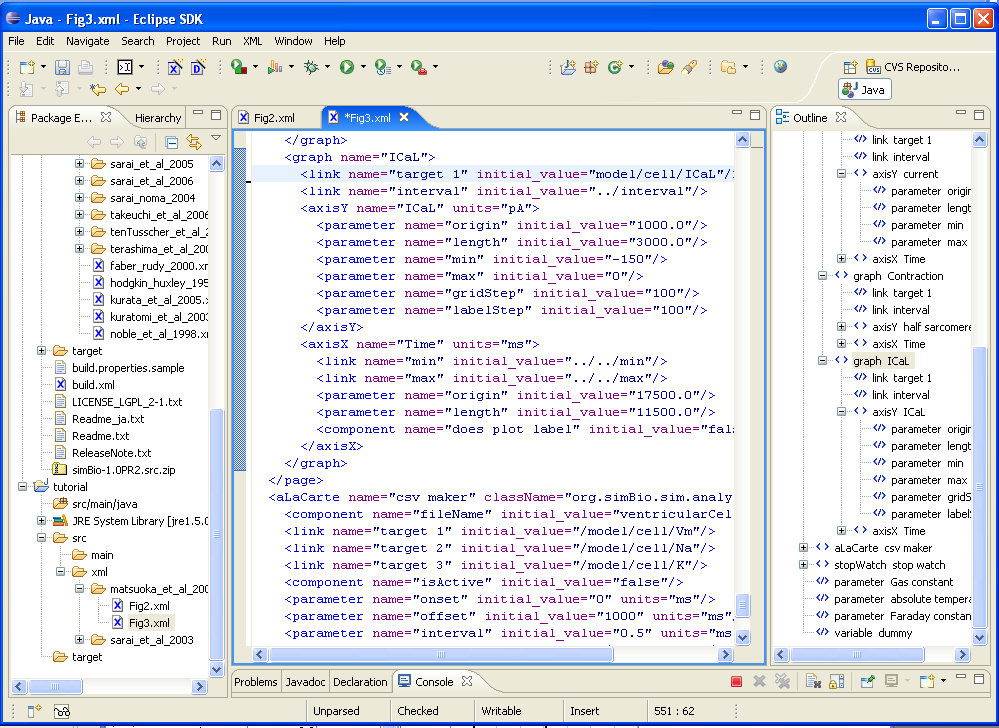

### 3-1-08.png

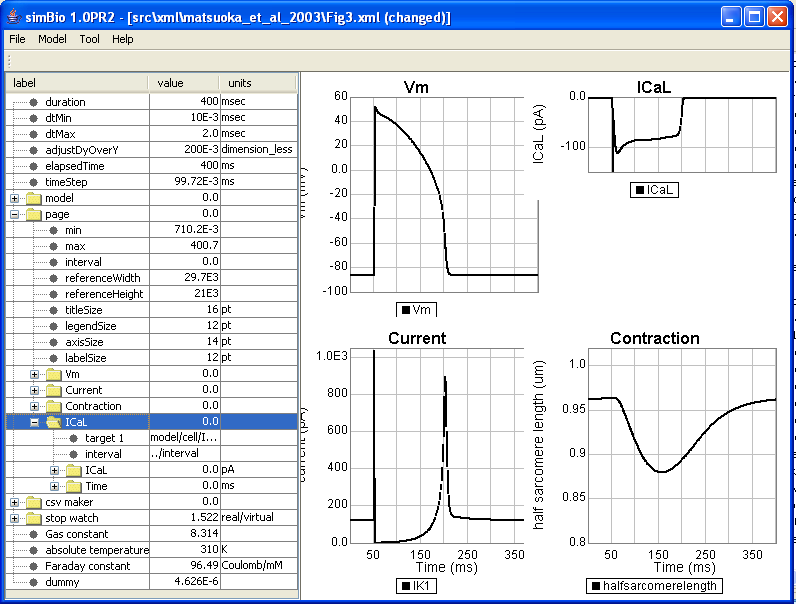

### 3-1-09.png

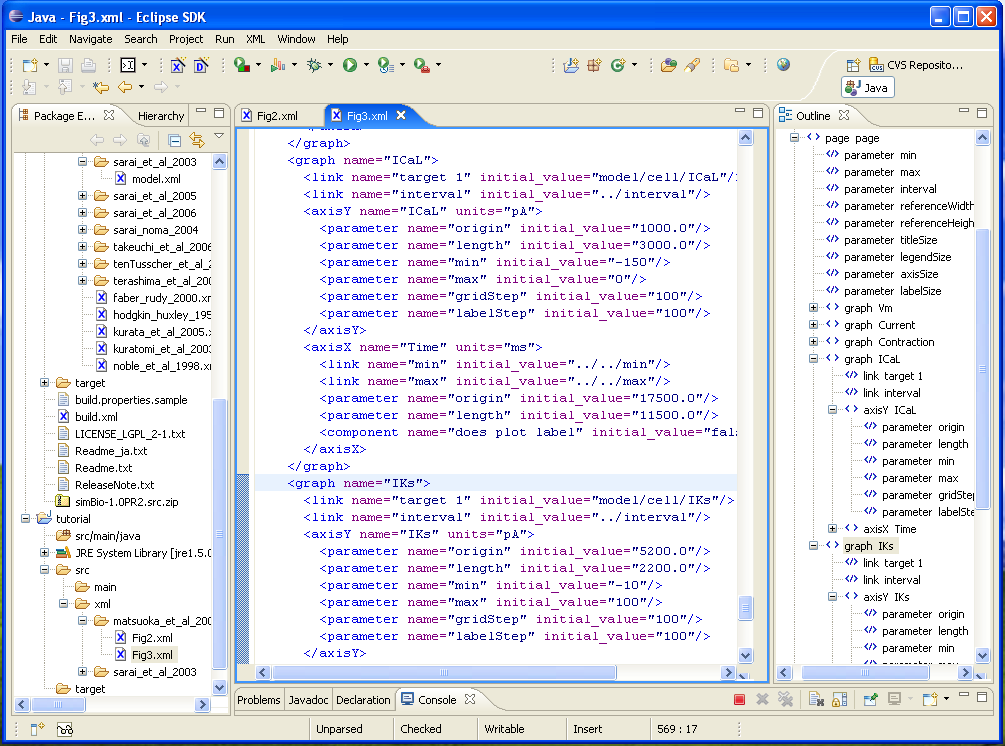

### 3-1-10.png

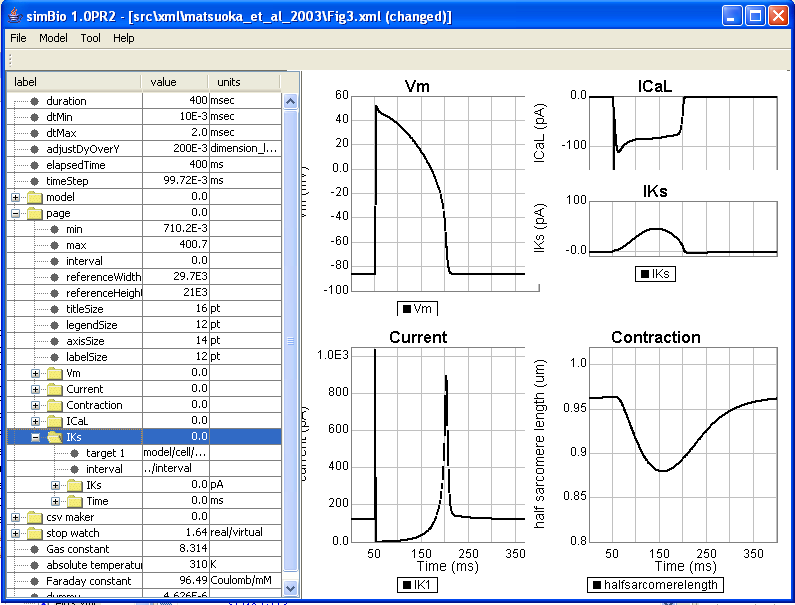

### 3-1-11.png

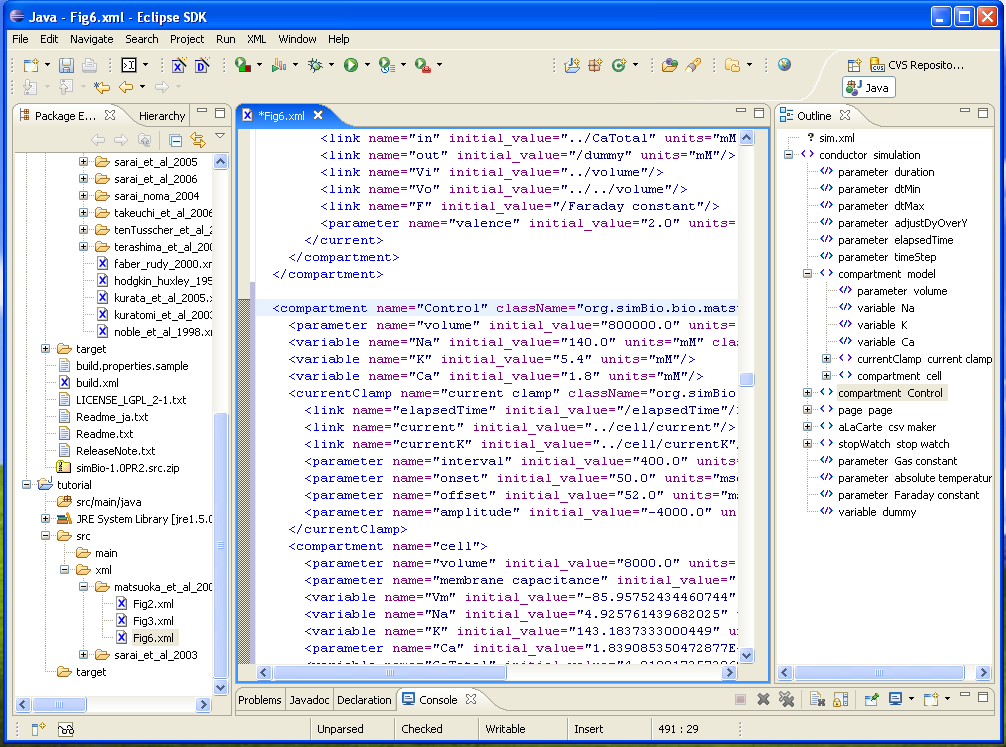

### 3-1-12_XML_ICaL.png

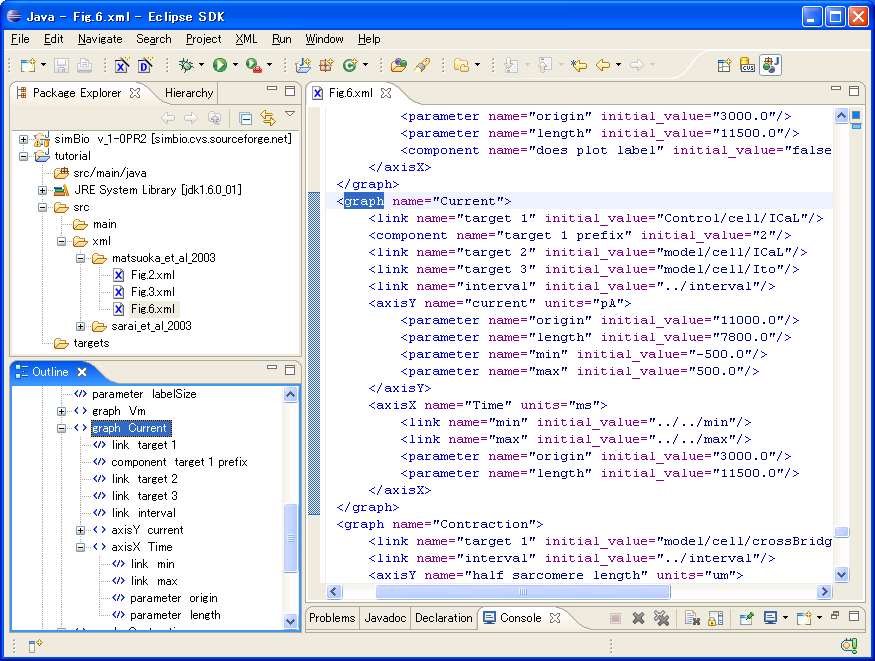

### 3-1-13_Figure6.png

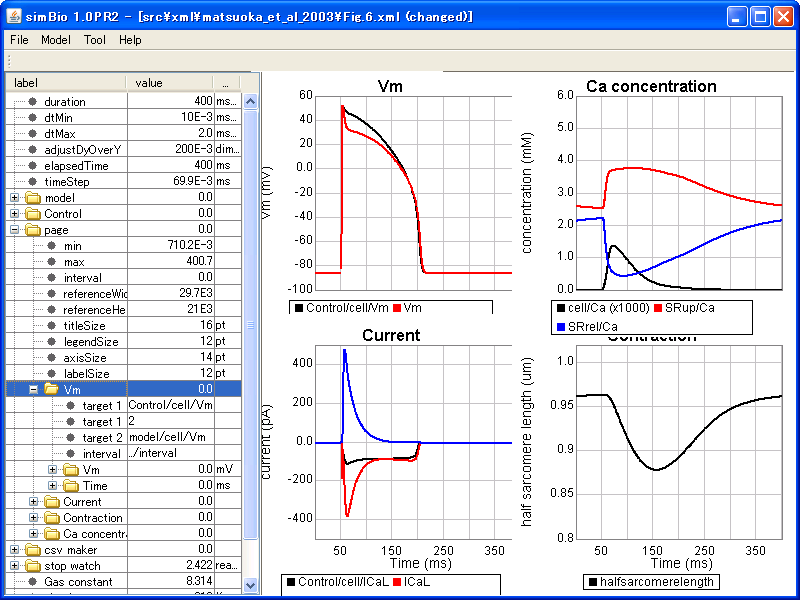

### 3-2-01.png

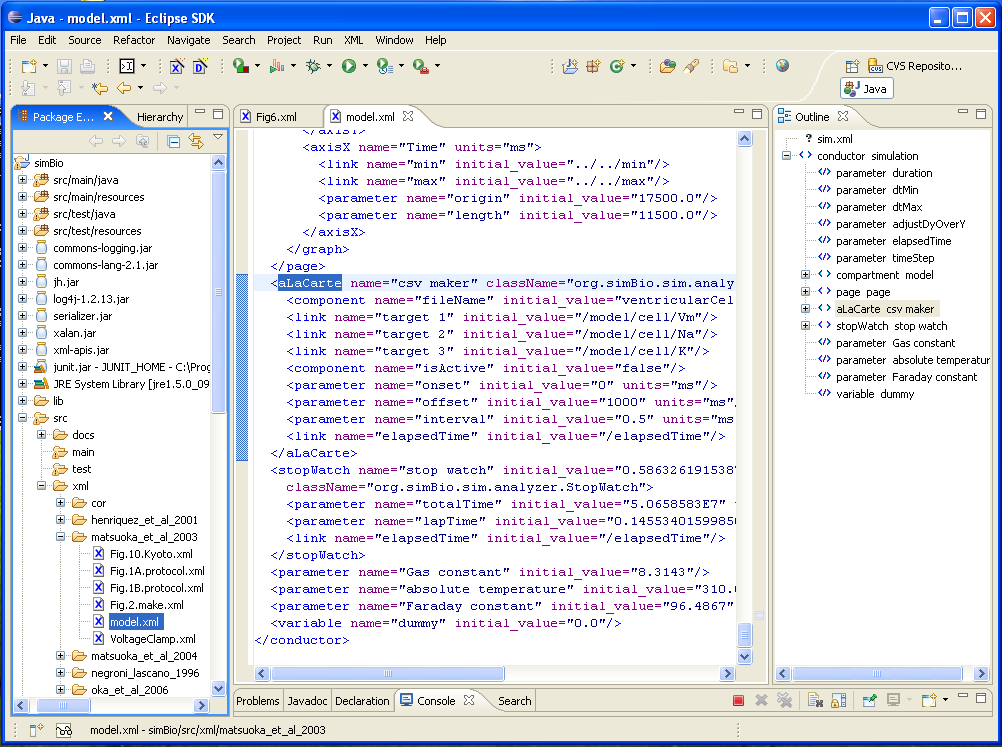

### 3-2-02.png

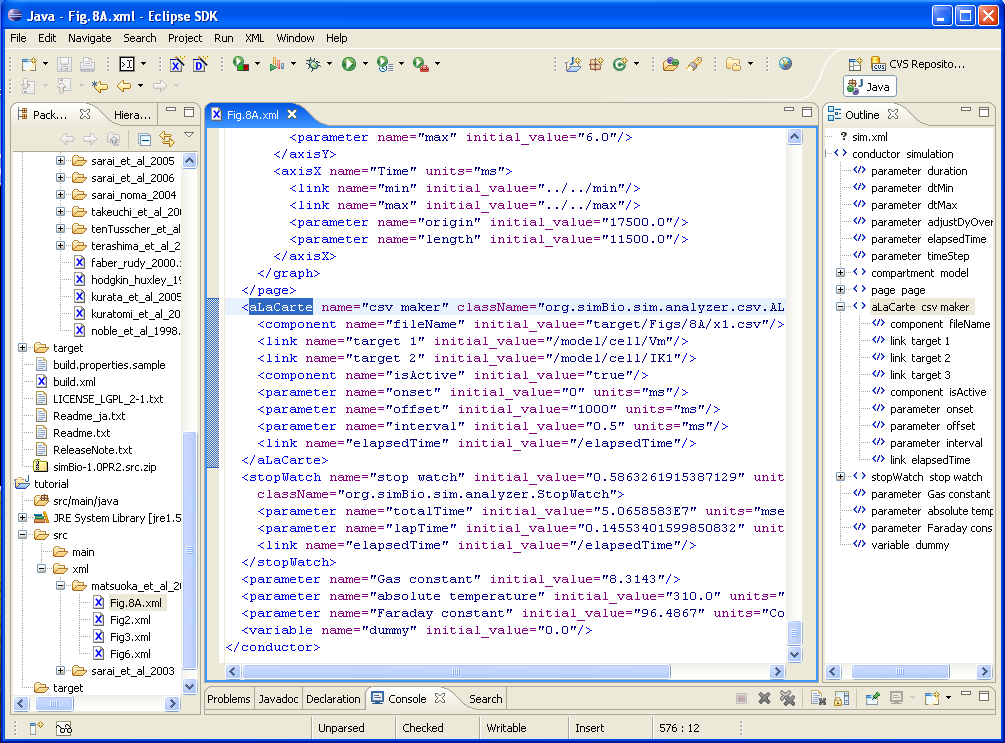

### 3-2-03_openCsv.png

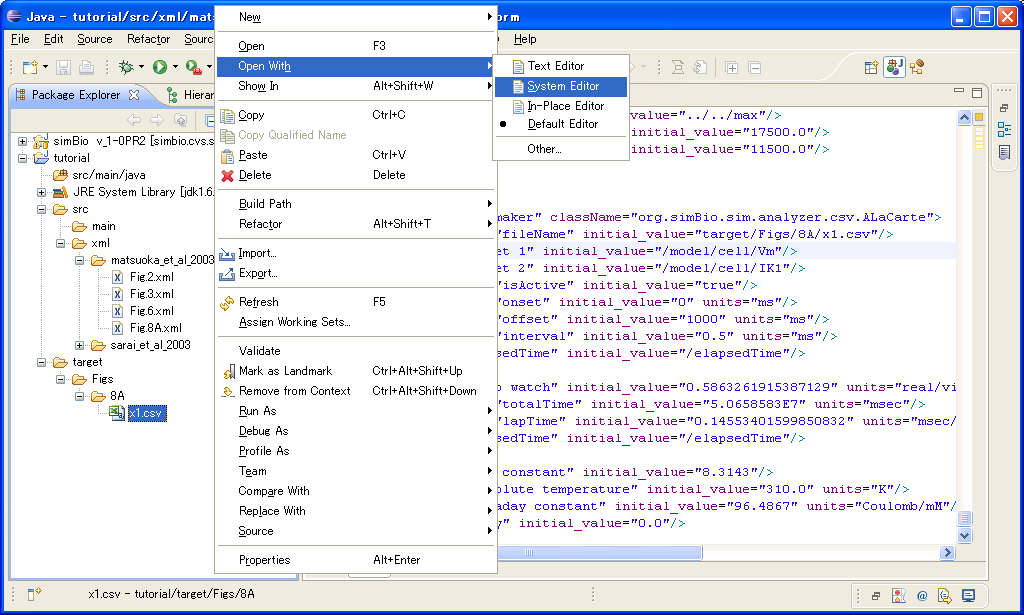

### 3-2-04.png

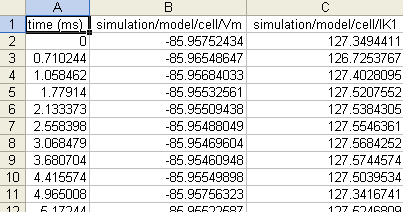

### 3-2-05.png

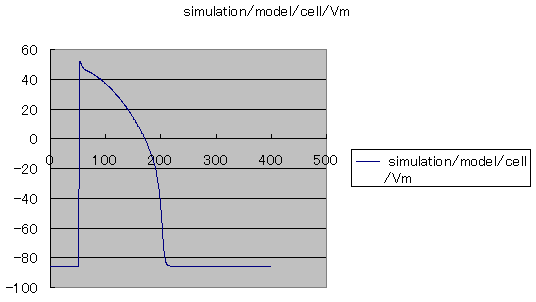

### 3-2-06.png

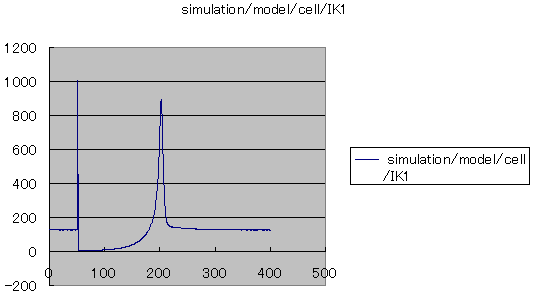

### 3-3-01_mini.png

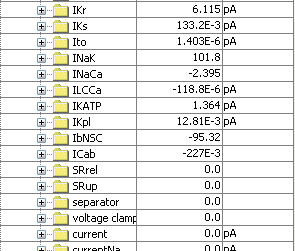

### 3-3-02.png

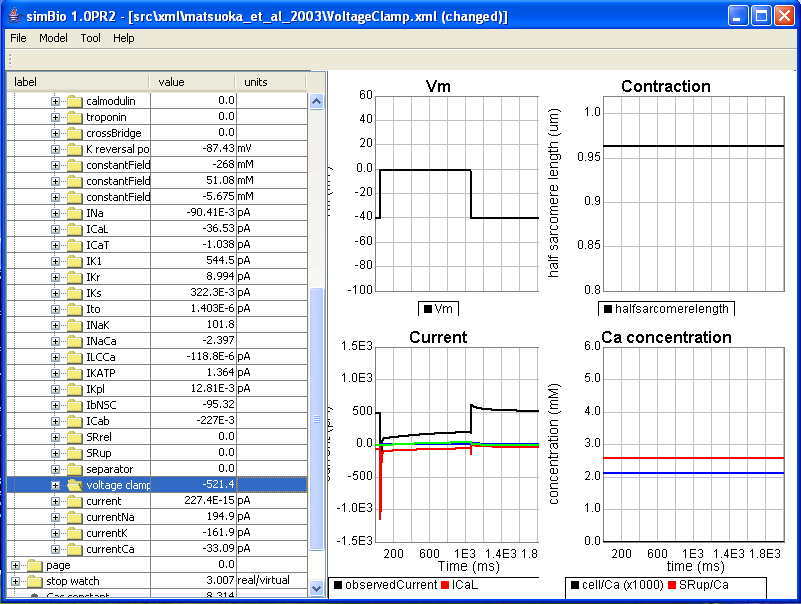

### 3-3-03_exchanger.png

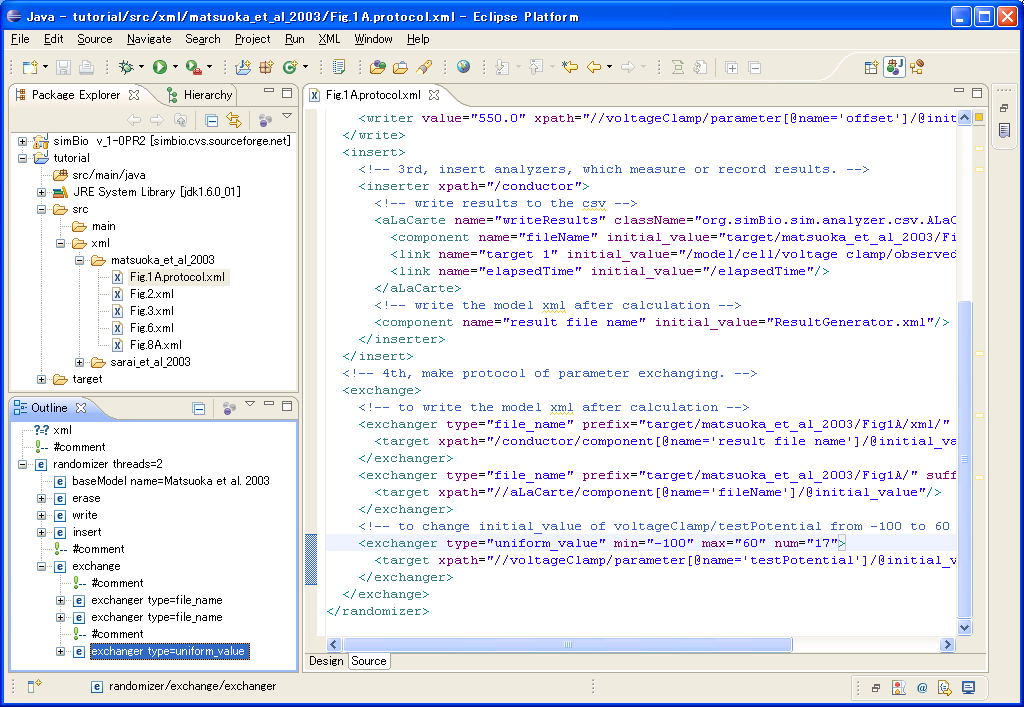

### 3-3-04_RunProtocol.png

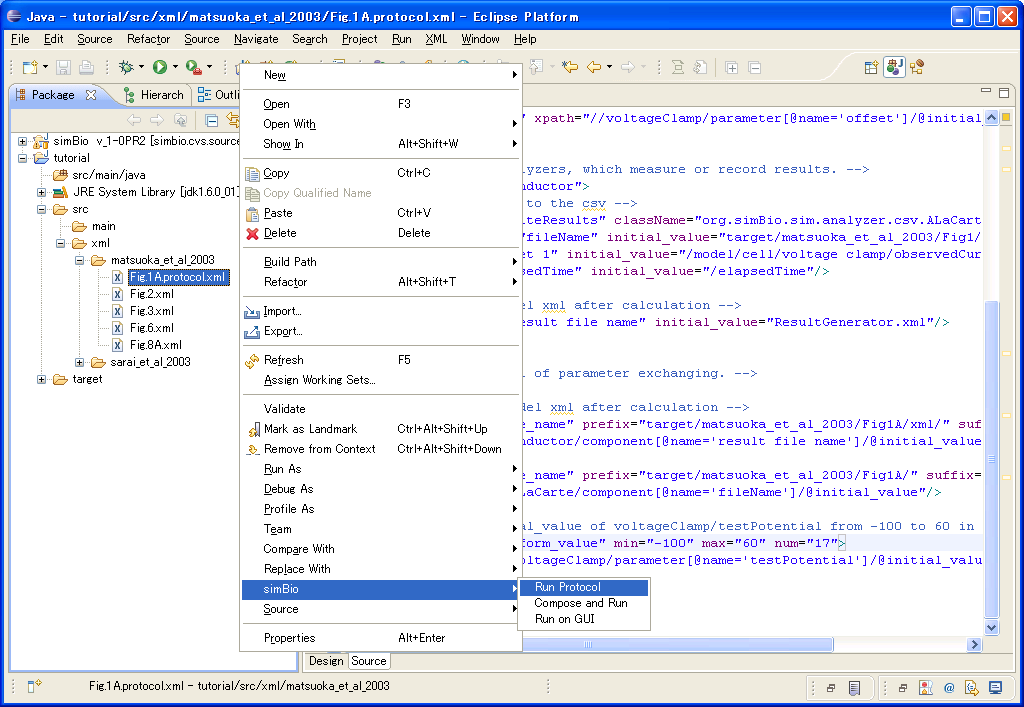

### 3-3-05_CSVs.png

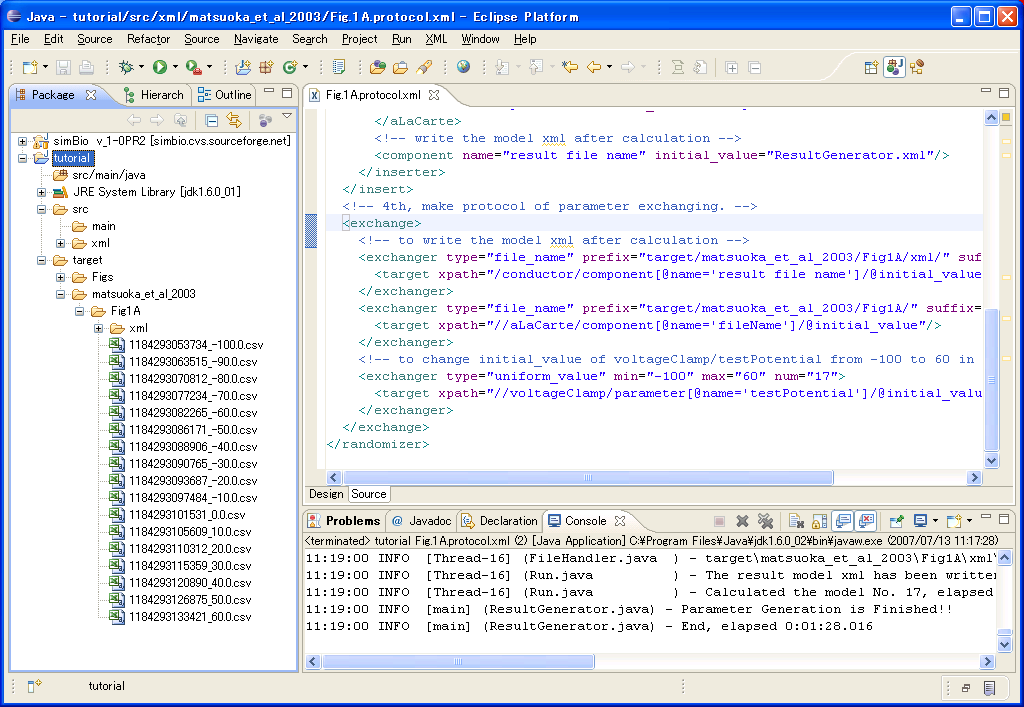

### 3-3-06_0to60.png

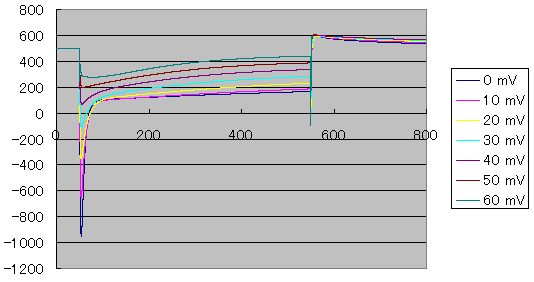

### 3-3-07.png

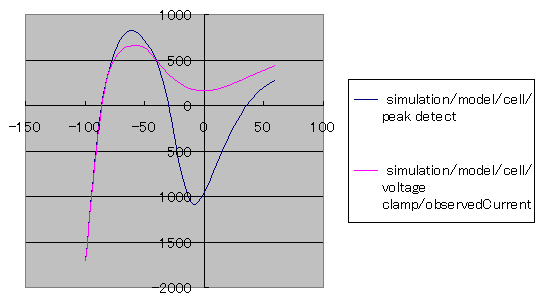

### 3-4-01_Fig.1A.protocol.xml.png

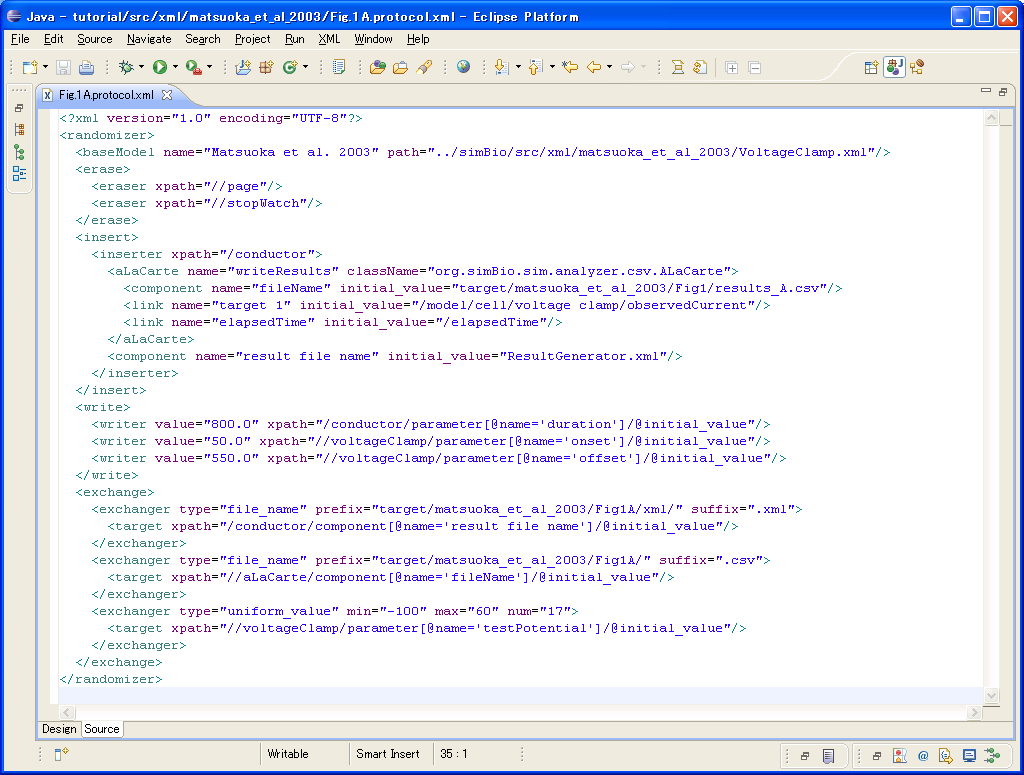

### 3-4-02.png

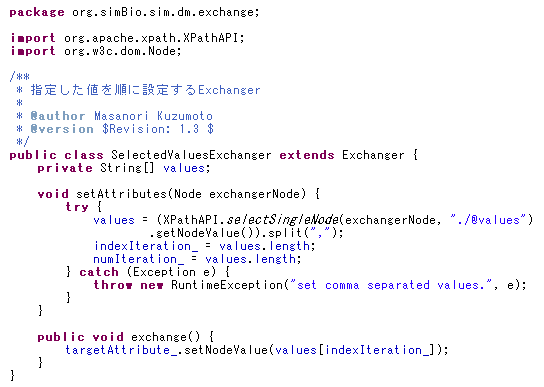

### 3-4-03.png

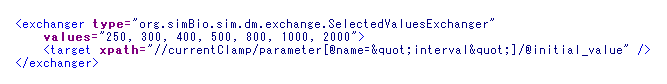

### 3-4-04.png

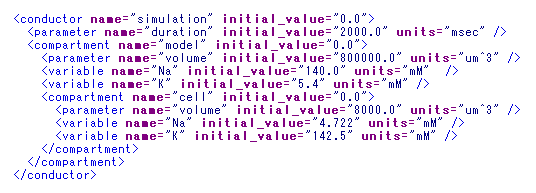

### 3-5-01.png

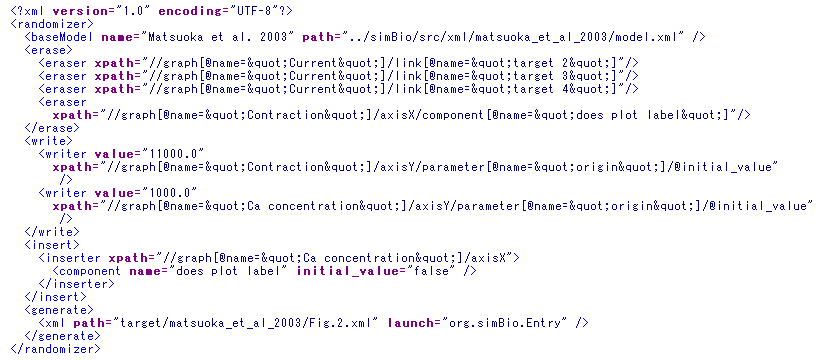

### 4-5-01_generateXML.png

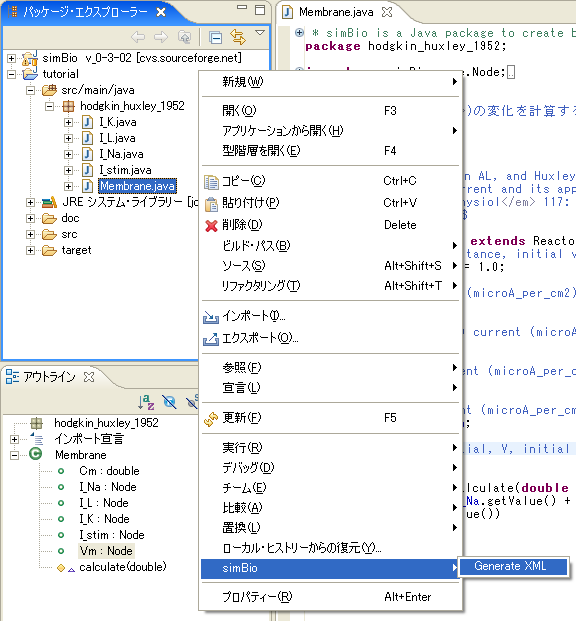

### ActivehttpServer.png

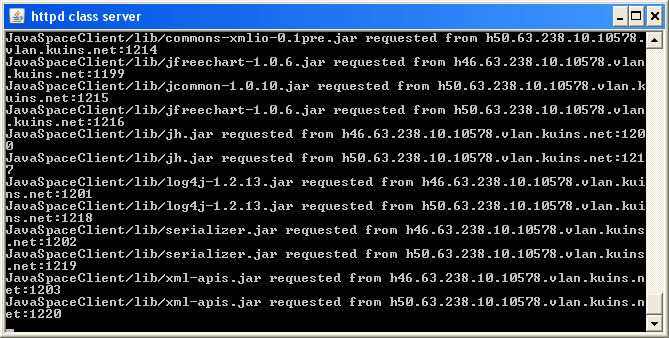

### Benchmark.png

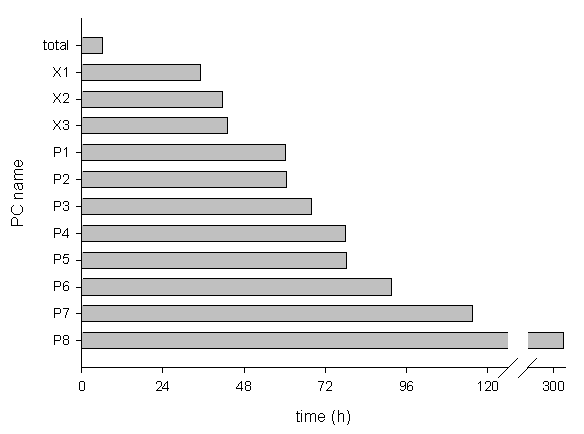

### CommandServer.png

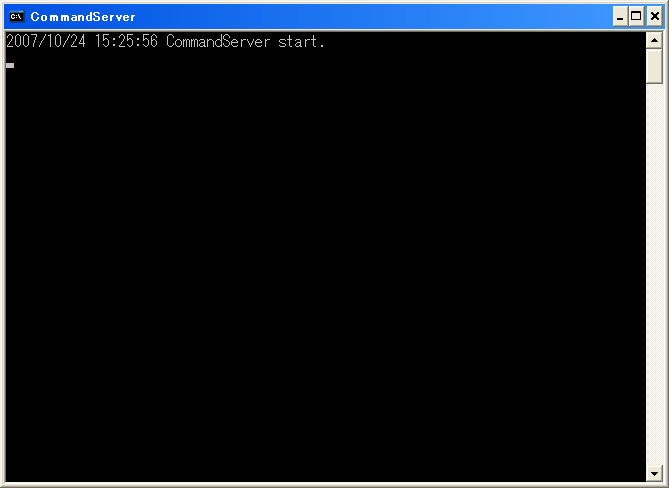

### DistributedSystem.png

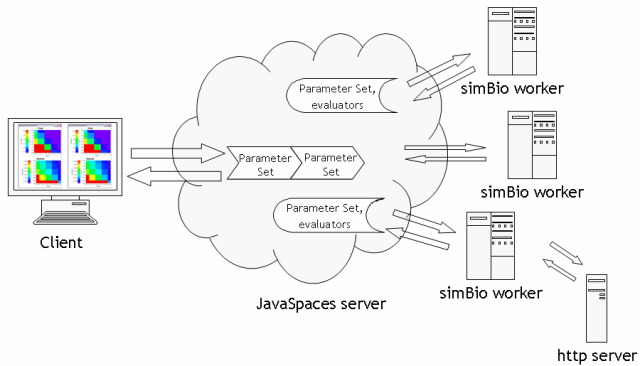

### group-logo.gif

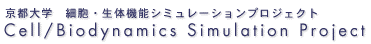

### httpServer.png

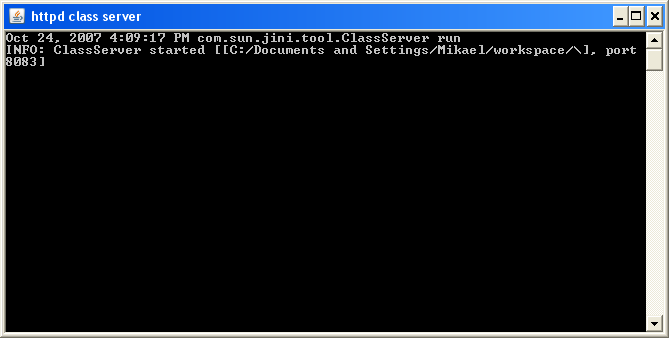

### JavaSpacesServer.png

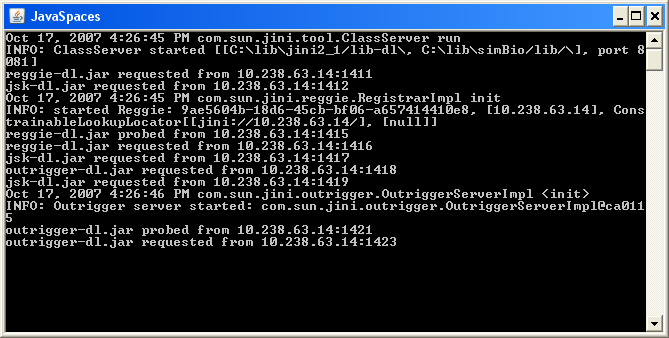

### modelDriven.png

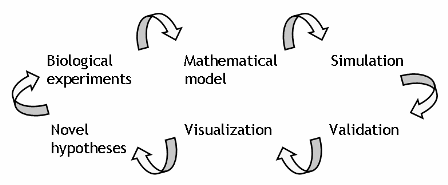
